# The histone methyltransferase DOT1L is essential for humoral immune responses

**DOI:** 10.1101/821462

**Authors:** Liam Kealy, Andrea Di Pietro, Lauren Hailes, Sebastian Scheer, Lennard Dalit, Joanna R Groom, Colby Zaph, Kim L Good-Jacobson

**Affiliations:** Department of Biochemistry and Molecular Biology, Monash University, Clayton, Victoria 3800, Australia; Infection and Immunity Program, Biomedicine Discovery Institute, Monash University, Clayton, Victoria 3800, Australia; Divisions of Immunology and Molecular Immunology, Walter and Eliza Hall Institute of Medical Research, Parkville, Victoria 3052, Australia; Department of Medical Biology, University of Melbourne, Parkville, Victoria 3010, Australia

**Keywords:** B cells, antibody, epigenetics, DOT1L, humoral responses, H3K79

## Abstract

Histone modifiers are essential molecular regulators that underpin the ability of immune cells to reprogram their gene expression during differentiation. The recruitment of the histone methyltransferase DOT1L induces oncogenic gene expression in a subset of B cell leukemia. Despite its importance, little is known about its role in the humoral immune system. Herein, we demonstrate that DOT1L is a critical regulator of B cell biology. *Dot1l*^*f/f*^*Mb1*^Cre/+^ mice had a block in B cell development, culminating in a significant reduction of mature B cells in the periphery. Upon immunization or influenza infection of *Dot1l*^*f/f*^*Cd23*^Cre/+^ mice, germinal centers failed to form and class-switched antibody-secreting cells were significantly attenuated. Consequently, immunized mice revealed that DOT1L was essential for the formation of B cell memory populations. Transcriptome, pathway and histological analysis identified a key role for DOT1L in reprogramming gene expression for migration and localization during the initial stages of a humoral response. Together, these results demonstrate an essential role for DOT1L in antigen-dependent B cell differentiation and hence, in generating an effective and lasting humoral immune response.

## INTRODUCTION

A successful antibody-mediated immune response leads to pathogen clearance and the formation of immune memory. These processes underpin the vast majority of vaccines (Good-Jacobson, 2018). The foundation for this success originates from the activation of a small number of antigen-specific naïve B cells. After activation, B cell differentiation is instructed by specific molecular programs that regulate fate and function during an immune response. These programs are modulated by epigenetic regulators (e.g. histone modifiers, siRNAs, DNA methylases) and transcription factors working in concert to determine whether genes are activated or repressed (Zhang and Good-Jacobson, 2019). Histone modifiers regulate chromatin accessibility, thereby regulating the accessibility of transcriptional machinery to their targets (Strahl and Allis, 2000). Importantly, the same histone modifiers emerging as important regulators of B cell differentiation can also become dysregulated in malignancies (Alberghini et al., 2015; Beguelin et al., 2013; Inaba et al., 2013). Thus, histone modifiers are therapeutically attractive to target with small molecule inhibitors (Arrowsmith et al., 2012). It is therefore important to understand the roles of histone modifiers in regulating B cell differentiation in both primary and secondary lymphoid organs.

DOT1L (*Disruptor of Telomeric Silencing 1-Like*) is the sole known enzyme that methylates lysine 79 of histone H3 (H3K79), and is most well-known for its role in oncogenesis. Approximately 40% of pediatric leukemias have been linked to H3K79 methylation-mediated oncogenic gene expression (Worden et al 2019). B cell Acute Lymphoblastic Leukemia (B-ALL) is an aggressive malignancy that affects children and adults, with long-term survival rates of <40% (Geng et al., 2012). A prominent subtype is *MLL* (*Mixed Lineage Leukaemia*)-rearranged B-ALL (*MLL*r B-ALL), The defining feature of *MLL*r-ALLs is rearrangement of the epigenetic regulator *MLL*, which results in the binding of MLL with any one of over 50 fusion partners. While this promiscuous partnering makes it difficult to find genuine therapeutic targets, another prominent feature of this leukaemia is the recruitment of a second epigenetic regulator, DOT1L (Krivtsov et al., 2008; Okada et al., 2005). Malignant gene expression is driven by DOT1L activity: inactivation of DOT1L downregulates genes targeted by MLL-fusion protein complexes (Bernt et al., 2011). Thus, DOT1L inhibition has become a major focus of translational research and is now in phase I/II clinical trials. Despite the clear importance of this molecule, there is scant information on the role of DOT1L in lymphocyte development in primary lymphoid organs or differentiation during an immune response in the periphery.

Epigenetic regulation of B cell differentiation is a nascent field, with understanding mostly limited to the roles of the methyltransferases EZH2 (Beguelin et al., 2013; Caganova et al., 2013), MLL2 (Zhang et al., 2015), demethylase LSD1 (Good-Jacobson, 2019; Haines et al., 2018; Hatzi et al., 2019) and acetyltransferase MOZ (Good-Jacobson et al., 2014) in germinal center (GC) biology. DOT1L was recently found to be expressed by human tonsillar and lymph node GC B cells (Szablewski et al., 2018). Differential H3K79 methylation on the *IgH* locus in plasma cells compared to B cells has been hypothesized to mediate the transition from membrane-bound to secretory forms of immunoglobulin (Milcarek et al., 2011). Thus, given its prominent role in *MLL*r B-ALL, its expression by GC B cells and potential role in antibody production, we set out to investigate the function of DOT1L in B cell differentiation and function.

By generating novel mouse strains deficient in DOT1L in either developing B cells or mature B cells, we describe previously unknown functional roles for DOT1L in B cell differentiation in both primary and secondary lymphoid organs. Our experiments delineate DOT1L as a vital checkpoint molecule during an immune response, as evidenced by the complete abrogation of immune responses to a range of immunization and infection models. In particular, our study links histone modification to the regulation of a network of migration-related genes and ultimately, B cell positioning in the follicle. Together, these results reveal the absolute requirement for DOT1L for B cells to mount an effective immune response in vivo.

## RESULTS AND DISCUSSION

### DOT1L is required for B cell development

We first sought to determine whether DOT1L was required in B cell biology through the generation of *Dot1l*^*f/f*^*Mb1*^Cre/+^ mice in which exon 2 of *Dot1l* was excised upon *Igα* expression (Figure S1A). Spleens from *Dot1l*^*f/f*^*Mb1*^Cre/+^ mice were notably smaller than those from *Mb1*^Cre/+^ control mice (Figures S1B-S1C), and the B cell population was significantly reduced in *Dot1l*^*f/f*^*Mb1*^Cre/+^ mice compared to both *Dot1l^f/+^Mb1*^Cre/+^ and *Mb1*^Cre/+^ controls (Figure S1D). Histological analyses revealed DOT1L-deficient mice were able to form follicular structures, however a thinner marginal zone structure was clearly evident (Figure S1E). Correspondingly, both splenic marginal zone and follicular B cells were significantly decreased (Figures S1F-S1I). Furthermore, circulating antibodies of all isotypes were significantly decreased in the absence of DOT1L (Figures S1J-S1O).

Thus, it appeared that DOT1L was required for efficient production of B cells to populate secondary lymphoid organs such as the spleen. To investigate, the production of developing B cell subsets in the bone marrow of *Dot1l*^*f/f*^*Mb1*^Cre/+^ and *Dot1l^f/+^Mb1*^Cre/+^ were compared to *Mb1*^Cre/+^ controls. Deficiency in DOT1L resulted in a clear decrease in the B cell population (Figure S1P). In keeping with the deletion of *Dot1l* at the pro-B cell stage in this conditional deletion, there was no significant change in either pre-pro B cells (data not shown) or pro-B cells (Figure S1Q). There was, however, a ~3-fold significant decrease in both the frequency (data not shown) and number (Figures S1Q) in all subsequent developing B cell subsets. Furthermore, in most instances, mice heterozygous for the floxed allele suggested a gene dosage effect of DOT1L during the development of B cells (Figure S1). These data demonstrate an essential requirement for DOT1L during B cell development in the bone marrow and thus establishment of peripheral B cell populations.

### DOT1L is essential for formation of GCs

DOT1L was recently found to be expressed in a subset of human GC B cells, suggesting a role for DOT1L in the GC (Szablewski et al., 2018). The significantly reduced number of peripheral B cells in *Dot1l*^*f/f*^*Mb1*^Cre/+^ mice made the precise role of DOT1L in antigen-driven responses difficult to disentangle from the developmental defects in these mice. Therefore, in order to determine the role of DOT1L during B cell responses, we generated novel mice in which *Dot1l^f/f^* mice were crossed with *Cd23*^cre^ mice. These mice deleted *Dot1l* specifically in mature B cells. Thus, any phenotype in *Dot1l*^*f/f*^*Cd23*^Cre/+^ mice should specifically be a consequence of the role of DOT1L in peripheral B cell differentiation and not due to the role of DOT1L observed during B cell development in the bone marrow (Fig. S2A-B). *Cd23*^cre/+^ and *Dot1l*^*f/f*^*Cd23*^Cre/+^ mice were immunized with the hapten (4-Hydroxy-3-nitrophenyl)-acetyl conjugated to Keyhole Limpet Hemocyanin (NP-KLH) precipitated on the adjuvant alum and GC B cell responses assessed at d7 and d28. At day 7, control mice generate a discernible NP^+^CD95^hi^ GC B cell population (Figure 1A). In contrast, conditional deletion of DOT1L in B cells prohibited the formation of GC (Figure 1A). This was not due to delayed kinetics. Strikingly, GC B cell frequency (Figure 1B) and number (Figure 1C) were completely absent in *Dot1l*^*f/f*^*Cd23*^Cre/+^ mice assessed during early (day 7) and late (day 28) phases of the response. While splenic morphology of B cell follicles appeared normal in *Dot1l*^*f/f*^*Cd23*^Cre/+^ mice compared to *Cd23*^cre/+^ controls (Figures 1D-1E), histological analyses showed a complete lack of PNA^+^ GC (Figure 1D) and IgG1^+^ cells (Figure 1E). Therefore, DOT1L was essential for the formation of GC and IgG1^+^ B cells during an immune response against alum-precipitated NP-KLH.

**Figure 1:**
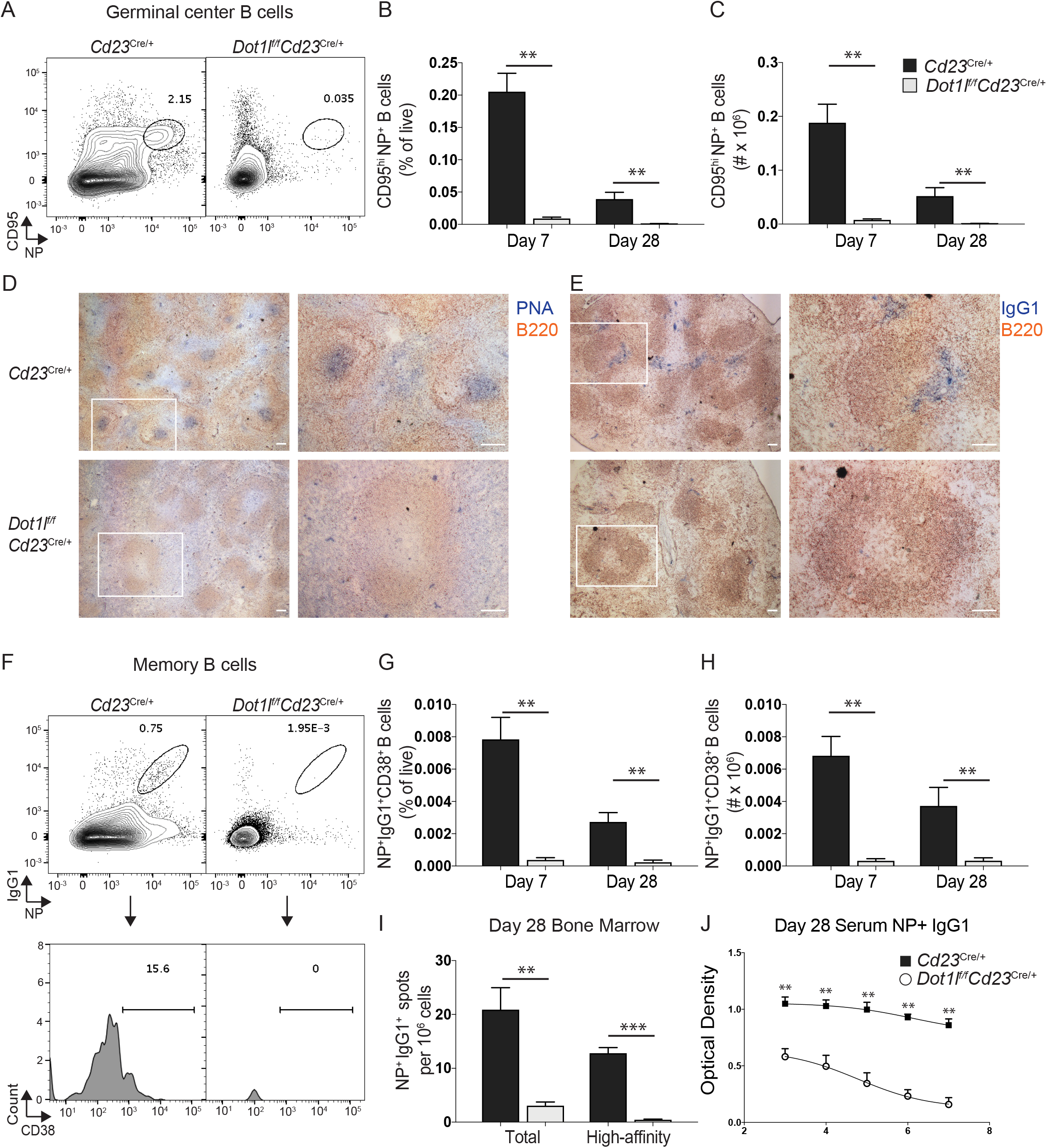
Dot1l deletion results in ablation of GCs and inability to form humoral memory. (A) Flow cytometric representative plot of GC B cells in *Cd23*^Cre/+^ and *Dot1l*^*f/f*^*Cd23*^Cre/+^ mice 7 days post-immunization with NP-KLH precipitated in alum. (B) *Cd23*^Cre/+^ (black bars or closed squares) and *Dot1l*^*f/f*^*Cd23*^Cre/+^ (white bars or open circles) mice were immunized with NP-KLH precipitated in alum and assessed at days 7 and 28 post-immunization for (B) frequency and (C) number of splenic CD19^+^IgD^−^NP^+^CD95^+^ cells. (D) Representative images from histological analyses day 7 post-immunization: B220 (red) and PNA (blue). Scale bar = 100μm. (E) Representative images from histological analyses day 7 post-immunization: B220 (red) and IgG1 (blue). Scale bar = 100μm. (F) Flow cytometric representative plot of NP^+^IgG1^+^CD38^+^ memory B cells. (G) Frequency and (H) total number of memory B cells in the spleen. (I) ELISpot analysis of IgG1^+^NP^+^ ASCs in the bone marrow at day 28 post-immunization. (J) ELISA analysis of NP^+^IgG1 serum antibody 28 days post-immunization. Results are compiled from experiments performed at two separate time points: day 7 (*Cd23^Cre/+^* (n=6) and *Dot1l^f/f^Cd23^Cre/+^* (n=6) mice; combined from two independent experiments) and day 28 (n=7 per genotype; combined from two independent experiments) post-immunization with NP-KLH in alum. ** P < 0.01; *** P < 0.001.

### DOT1L is essential for the establishment of humoral immunity

The formation of immune memory is a critical outcome for an effective B cell response. Immune memory populations in the humoral arm of the immune system consist of memory B cells and long-lived plasma cells. In several gene-deficient models, the abrogation of GC responses is associated with an increase in early memory B cells (Good-Jacobson and Shlomchik, 2010). We therefore determined whether the absence of DOT1L affected the formation of immune memory subsets. *Cd23*^cre/+^ and *Dot1l*^*f/f*^*Cd23*^Cre/+^ mice were immunized with NP-KLH in alum and assessed at both day 7 and 28 post-immunization. Memory B cells, as defined by NP^+^IgG1^+^CD38^+^, were completely absent in DOT1L-deficient mice (Figure 1F). At both days 7 and 28 post-immunization, memory B cell frequency (Figure 1G) and number (Figure 1H) were absent in *Dot1l*^*f/f*^*Cd23*^Cre/+^ mice. Correspondingly, plasma cells in the bone marrow (Figure 1I) were also significantly decreased, with high-affinity antibody-secreting cells (ASCs) completely absent. As such, circulating NP-binding IgG1 antibody was significantly decreased (Figure 1J). Therefore, DOT1L was essential for the establishment of humoral memory following T-dependent immunization.

### DOT1L is required for GC B cells in response to the Th1 cell-biased influenza infection

Inhibition of DOT1L in T helper (Th) cells was shown to result in an increase in both Th1 cells and IFNγ production by those cells in vitro (Scheer et al., 2019). B cells also have tailored functional responses to either Th1 cell-biased or Th2 cell-biased stimuli, and although transcription factor expression is associated with dichotomous responses (Piovesan et al., 2017), it is not known whether epigenetic regulators are also important in specialized B cell function. Given the modulation of Th cell function by DOT1L, we used an influenza model to assess the role of DOT1L in an IFNγ-mediated B cell response and determine whether the role of DOT1L is generalizable across different types of responses in B cells. *Cd23*^cre/+^ and *Dot1l*^*f/f*^*Cd23*^Cre/+^ mice were infected with the H3N2 HKx31 influenza virus. B cell differentiation was assessed in the spleen and draining (mediastinal) lymph node 8 and 14 days post-infection. Concordant with the immunization results (Figure 1), DOT1L was required for B cells to mount an effective response to influenza. In the absence of DOT1L, there was a ≥5-fold decrease in splenic GC B cells (Figures S2C-2E) and a similar reduction of GC B cells in the mediastinal lymph node (Figures S2F-2G). Those GC B cells that remained were examined for isotype switching to IgG2c, and found to be ~2-fold decreased on a per GC B cell basis (Figure S2H). Finally, we excluded aberrant skewing of immunoglobulin isotypes by assessing Th2-associated IgG1 production in influenza-infected (Th1 cell-biased) mice, and conversely, Th1-associated IgG2c production in NP-KLH-immunized (Th2 cell-biased) mice. In both cases, antibody production of both isotypes was significantly reduced in the absence of DOT1L and thus there was no evidence of aberrant Th cell-biased responses these mice (Figures S2I and S2J, respectively). Therefore, DOT1L was essential for B cells to mount an effective GC reaction during either Th1 or Th2 cell-biased immune responses.

### DOT1L is required for isotype-switched plasmablast formation in vivo

During the early phases of a T-dependent immunization, activated B cells either migrate back into the follicles and form GC B cells, or they can form foci of proliferating extrafollicular plasmablasts secreting low-affinity unswitched (IgM) and switched (IgG1) isotypes. As GC were unable to form in the absence of DOT1L, we asked whether this was specific to GC B cell differentiation, or whether plasmablast differentiation was also abrogated in vivo. We first examined the frequency of formation of extrafollicular plasmablasts at day 7 post immunization. *Cd23*^cre/+^ and *Dot1l*^*f/f*^*Cd23*^Cre/+^ mice were immunized with NP-KLH in alum and splenic plasmablasts assessed as B220^lo^CD138^+^ cells (Figure 2A). In the absence of DOT1L, there was a 3-fold reduction in plasmablasts (Figures 2B-2C). While a small population of plasmablasts remained in *Dot1l*^*f/f*^*Cd23*^Cre/+^ mice, these cells had not switched to IgG1 (Figures 2D-2E). The formation of antigen-specific antibody-secreting plasmablasts, as well as circulating antibody, were also significantly diminished in the absence of DOT1L. NP-specific IgG1^+^ ASCs were absent from *Dot1l*^*f/f*^*Cd23*^Cre/+^ mice (Figures 2F-2G). Accordingly, NP-binding IgG1 antibody was not detected in serum (Figure 2H). In contrast, there was little difference in NP-specific IgM ASCs (Figure 2I) and only a small reduction in NP-binding serum IgM (Figure 2J). While this was consistent with a defect in isotype switching, it may instead reflect the timing of DOT1L function. That is, the defect observed in DOT1L-deficient mice may have occurred after the formation of IgM^+^ plasmablasts in this model. Lastly, we also assessed the formation of plasmablasts during an influenza infection. Plasmablasts in both spleen (Figures 2K-2L) and mediastinal lymph node (Figures 2M-2N), as well as serum IgG2c (Figure 2O), were significantly reduced. Together, these data demonstrated an absolute requirement for DOT1L for GC B cell and isotype-switched plasmablast formation in both Th2 and Th1 cell-biased responses.

**Figure 2:**
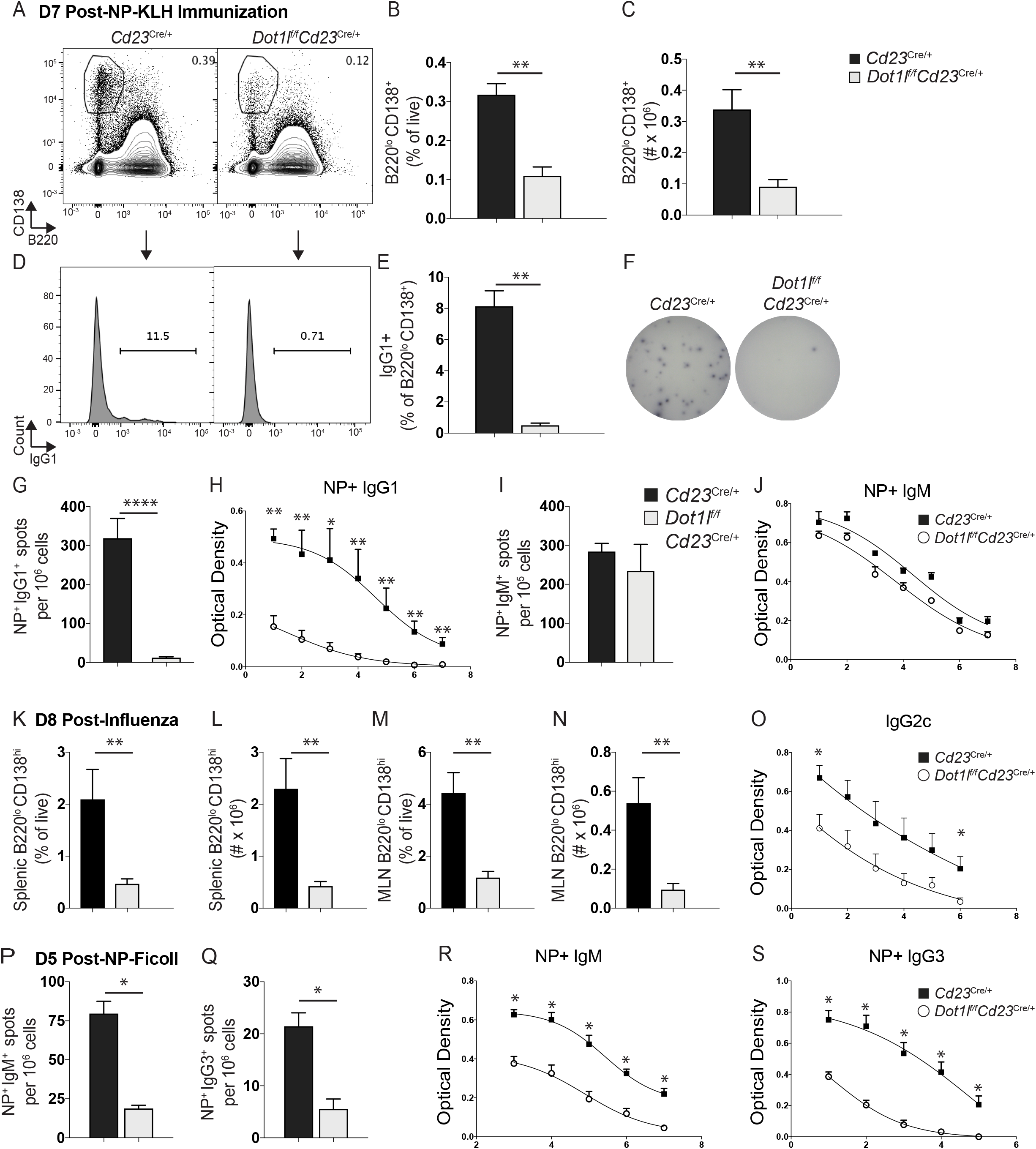
DOT1L plays a key role in the formation and class-switching of early splenic plasma cells produced in response to TD antigens. (A) Flow cytometry representative plot of B220^lo^CD138^+^ cells in the presence or absence of Dot1L 7 days post-immunization with NP-KLH in alum. (B-C) Frequency (B) and (C) number of B220^lo^CD138^+^ cells in *Cd23*^Cre/+^ (black bars) and *Dot1l*^*f/f*^*Cd23*^Cre/+^ mice (light grey bars). (D) Flow representative plot and (E) frequency of IgG1^+^ cells within the B220^lo^CD138^+^ population. (F-G) ELISpot analysis of NP^+^ IgG1^+^ ASCs in the spleen at day 7 post-immunization. (H) Serum NP^+^IgG1 antibody at day 7 post-immunization. (I) ELISpot analysis of NP^+^IgM^+^ ASCs in the spleen at day 7 post-immunization. (J) Serum NP^+^IgM antibody at day 7 post-immunization. Results are compiled from experiments performed at two separate time points: day 7 (*Cd23^Cre/+^* (n=6) and *Dot1l^f/f^Cd23^Cre/+^* (n=6) mice; combined from two independent experiments) and day 28 (n=7 per genotype; combined from two independent experiments) post-immunization with NP-KLH in alum. (K-N) *Cd23*^Cre/+^ and *Dot1l*^*f/f*^*Cd23*^Cre/+^ mice were infected with HKx31 influenza virus and assessed 8 and 14 days post-infection for (K) frequency and (L) number of B220^lo^CD138^+^ cells in the spleen, and (M) frequency and (N) number of B220^lo^CD138^+^ cells in the mediastinal lymph node. (O) Total serum IgG2c at day 14 post-infection. n=6 per genotype, combined from two independent experiments per time point. * P < 0.05; ** P < 0.01; **** P < 0.0001. (P) ELISpot analysis of splenic NP^+^IgM^+^ ASCs 5 days post-immunization with NP-Ficoll in *Cd23*^Cre/+^ (black bars or closed squares) and *Dot1l*^*f/f*^*Cd23*^Cre/+^ mice (light grey bars or open circles). (Q) Splenic NP^+^IgG3^+^ ASCs. (R) NP^+^ IgM and (S) NP^+^ IgG3 serum antibody. n=6 per genotype, combined from two independent experiments per time point. * P < 0.05.

### DOT1L is required for mounting a B cell response independent of T cells or GCs

The results generated thus far revealed that DOT1L activity was important for establishing a GC and plasmablast response post-immunization with a T-dependent antigen. However, it remained unclear if DOT1L was required for B cell responses that formed independently of T cell help or GCs, or possibly promoted the T-independent plasmablast pathway. To investigate these questions in more detail, *Cd23*^cre/+^ and *Dot1l*^*f/f*^*Cd23*^Cre/+^ mice were immunized with NP-Ficoll in PBS, we utilized a type 2 T-independent antigen and B cell differentiation assessed 5 days later. Both antigen-specific IgM^+^ (Figure 2P) and IgG3^+^ ASCs (Figures 2Q) were significantly reduced in the absence of DOT1L. Accordingly, circulating NP-binding IgM (Figure 2R) and IgG3 (Figure 2S) antibodies were both significantly reduced. Interestingly, there was a stronger defect in IgM in the T-independent response (Figure 32R) compared to the T-dependent response (Figure 2J). Nevertheless, these data demonstrate that DOT1L is required for effective B cell differentiation into ASCs and antibody production in both T-dependent and T-independent responses.

### DOT1L regulates expression of genes required for effective B cell responses in vivo

The next series of experiments were undertaken to delineate how DOT1L regulates B cell fate. Histone modifiers have been linked with cell cycle regulation of B cells (Beguelin et al., 2017; Caganova et al., 2013). Given the absence of B cell responses across multiple immunization and infection models, we utilized two separate approaches to determine whether DOT1L-deficient B cells were unable to be activated, and/or were prone to undergo apoptosis. In the first, we compared cell trace violet (CTV)-labeled B cells isolated from *Cd23*^cre/+^ and *Dot1l*^*f/f*^*Cd23*^Cre/+^ mice stimulated *in vitro* with CD40L, IL-4 and IL-5. In the second, we incubated stimulated CTV-labeled wild-type B cells with a small molecule inhibitor to DOT1L (Scheer et al., 2019). In both cases, proliferation was not significantly affected in the absence of DOT1L (Figures S2K and S2L). Thus, humoral responses were not prematurely aborted due to an intrinsic lack of cell proliferation in DOT1L-deficient B cells.

We next confirmed the target histone modification regulated by DOT1L in B cells. Specifically, we determined whether conditional deletion of DOT1L would result in a reduction of global H3K79me2 in B cell subsets (Figure S3A). As a control, we also tested polycomb repressive complex 2 (PRC2)-mediated H3K27me3, which has also been shown to be an important regulator of B cell biology (Beguelin et al., 2013; Caganova et al., 2013; Guo et al., 2018; Su et al., 2003). Splenic IgD^+^ and IgD^−^ B cells were sort-purified from immunized *Cd23*^cre/+^ and *Dot1l*^*f/f*^*Cd23*^Cre/+^ mice, together with *Eed^f/f^Cd23*^cre/+^ mice as controls for PRC2 function (Figure S3A; EED is the scaffolding protein required for PRC2 assembly). As expected, in the absence of DOT1L, there was a global decrease in H3K79me2 in B cell subsets, but not in the absence of EED (Figure S3B-S3C). Conversely, global H3K27me3 was reduced in B cells in the absence of EED, but not in the absence of DOT1L (Figures S3D-S3E). Thus, these results confirmed that DOT1L specifically mediates H3K79 methylation in B cells.

This lead to us to hypothesize that DOT1L-mediated H3K79 methylation regulated unique transcriptional programs following B cell activation, which we assessed by RNA-sequencing. CD19^+^IgD^+^ naïve B cells and CD19^+^IgD^−^ B cells were sort-purified from NP-KLH in alum-immunized mice, the latter population being enriched in activated B cells during an immune response. To enrich for the immediate gene regulatory events, without the compounded impact of the absence of GC and early memory B cell populations, we isolated B cell subsets 4 days post-immunization, a time at which there was little difference in the frequency of antigen-specific B cells in DOT1L-deficient compared to DOT1L-sufficient mice (data not shown). As expected, the greatest differences were observed between CD19^+^IgD^+^ (naïve) and CD19^+^IgD^−^ (activated) B cells, and then between control and DOT1L-deficient cells (Figure S4A). Of note, the majority of genes differentially expressed were upregulated in DOT1L-deficient CD19^+^IgD^+^ naïve B cells compared to control naïve B cells, with only 5 genes downregulated (Figures 3A-3B). In contrast, 30 genes were downregulated in DOT1L-deficient CD19^+^IgD^−^ B cells compared to controls (Figures 3C-3D), correlating to the association of H3K79me2 with actively transcribed genes during cell differentiation (Steger et al., 2008). There were 37 genes that overlapped between datasets (Figure S4B). Of these common genes, the vast majority (89%) were upregulated in the absence of DOT1L, with downregulated genes almost solely restricted to CD19^+^IgD^−^ B cells (Figures S4C, S4D and S4E, respectively).

**Figure 3:**
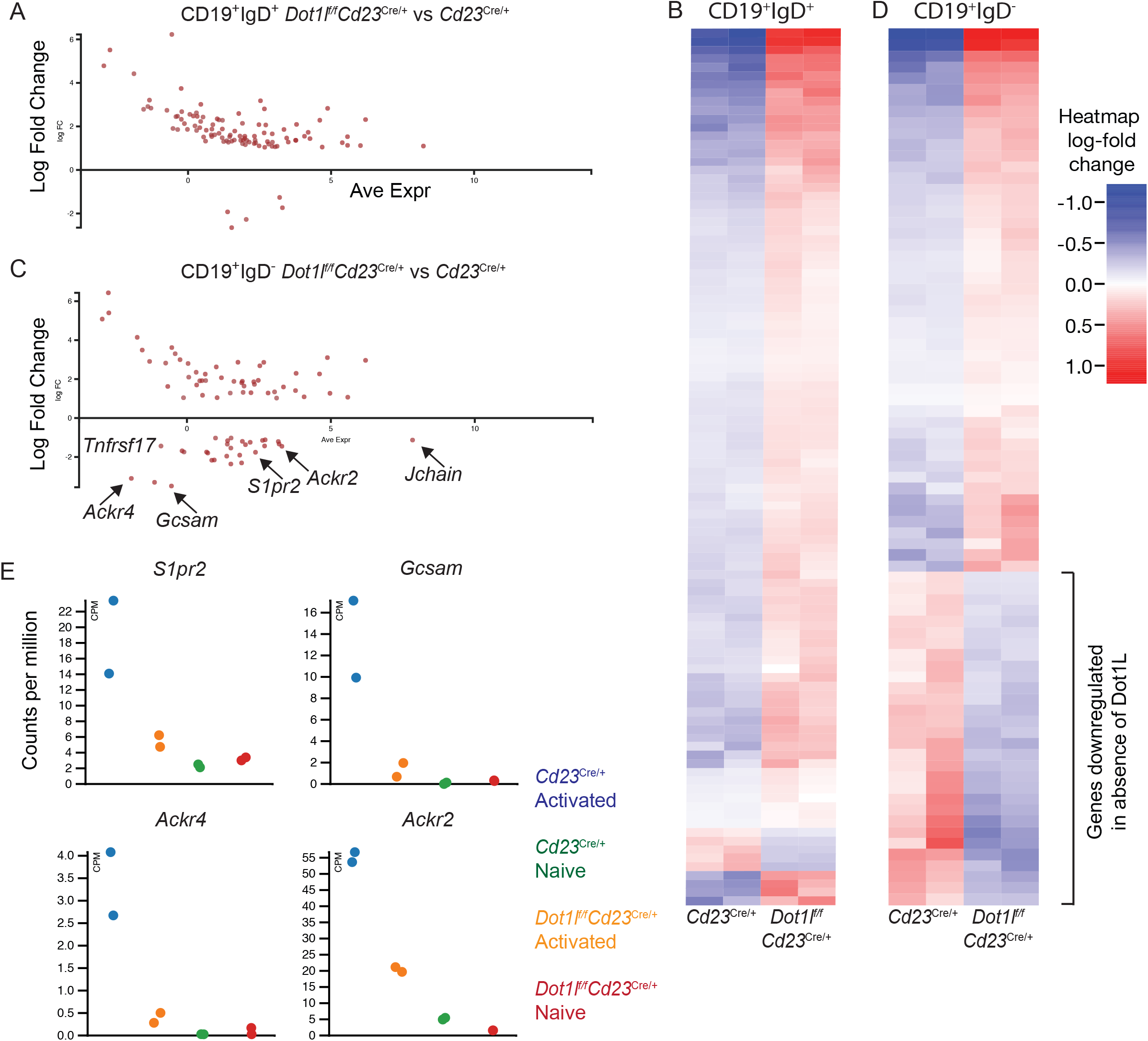
DOT1L regulates expression of genes required for effective B cell responses in vivo. (A-D) RNA-sequencing data: each sample is an independent sample obtained from either CD19^+^IgD^+^ or CD19^+^IgD^−^ B cells sort-purified from either *Dot1l*^*f/f*^*Cd23*^Cre/+^ or *Cd23*^Cre/+^ mice 4 days post-immunization with NP-KLH in alum. (A) MA plot for CD19^+^IgD^+^ samples. Shown is the average expression vs log fold change of *Dot1l*^*f/f*^*Cd23*^Cre/+^ samples compared to *Cd23*^Cre/+^ samples. A FDR cutoff of 0.01 and absolute log fold change of 1 was applied to the samples. (B) Heatmap of RNA-sequencing data (Table S1); each column is an independent sample obtained from sort-purified CD19^+^IgD^+^ B cells from either *Dot1l*^*f/f*^*Cd23*^Cre/+^ or *Cd23*^Cre/+^ mice immunized with NP-KLH in alum. (C) MA plot for CD19^+^IgD^−^ samples. Shown is the average expression vs log fold change of *Dot1l*^*f/f*^*Cd23*^Cre/+^ samples compared to *Cd23*^Cre/+^ samples. A FDR cutoff of 0.01 and absolute log fold change of 1 was applied to the samples. (D) Heatmap of RNAseq data (Table S2); each column is an independent sample obtained from sort-purified CD19^+^IgD^−^ B cells from either *Dot1l*^*f/f*^*Cd23*^Cre/+^ or *Cd23*^Cre/+^ mice immunized with NP-KLH in alum. (E) Plots showing count per million for individual genes for ‘naïve’ (CD19^+^IgD^+^) and ‘activated’ (CD19^+^IgD^−^) samples.

In other cell types, DOT1L function has been linked with inhibiting inappropriate differentiation (Wong et al., 2015). Thus, we interrogated whether the expression of transcription factors that regulate differentiation steps in B cells, such as *Bcl6* and *IRF4* (Dent et al., 1997; Klein et al., 2006; Willis et al., 2014), was dysregulated. However, there was no observable difference in expression of these transcription factors (data not shown). Multiple genes associated with B cell activation and migration, such as *Ackr2, Ackr4, GCsam, S1pr2* and *Tnfrsf17,* were upregulated in B cells from *Cd23*^cre/+^ upon activation, compared to naïve B cells, but were not expressed in either DOT1L-deficient subsets (Figures 3C-E). Chemokine receptors expressed on B cells regulate their ability to migrate in response to chemokine gradients present in the microenvironment (Lu and Cyster, 2019). The atypical chemokine receptors Ackr2 and Ackr4 has been linked to regulation of B cell migration (Hansell et al., 2011; Kara et al., 2018). While the role of Ackr2 in GC responses has not yet been studied, Ackr4 has recently been identified as a negative regulator of B cell responses with enforced expression of Ackr4 reducing B cell migration towards CCL21, a CCR7 ligand (Kara et al., 2018). Correspondingly, DOT1L-deficient cells, which had a reduction in *Ackr4*, had an increased ability to migrate to CCL21 (Figure S4F). Furthermore, S1PR2 (the G protein-coupled sphingosine-1-phosphate receptor encoded by *S1pr2*) is critical for B cell confinement in the GC (Green and Cyster, 2012). While we did not observe in increase in GCs observed in Ackr4-deficient (Kara et al., 2018) and S1PR2-deficient (Green and Cyster, 2012) mice, we hypothesized that disruption of the expression of multiple genes associated with early activation and migratory events in DOT1L-deficient B cells may result in early defective humoral responses in vivo.

### DOT1L regulates localization of activated B cells during a humoral response

During the first 3-4 days of an immune response, B cells migrate to the T:B border and the interfollicular zone before moving back into the follicle to establish GCs (Chan et al., 2009; Kerfoot et al., 2011; Kitano et al., 2011). To gain further insight into the networks of genes regulated by DOT1L during early B cell responses, Ingenuity Pathway Analysis (IPA) was performed. Notably, cellular movement, cell signalling and DNA replication and repair as the top three functions identified by IPA as being dysregulated in the absence of DO1L (Figure 4A). Further analysis revealed networks of molecules involved in cell migration (Figure 4B) and B cell responses (Figure S4G). To investigate the effect of changes in these molecules, we immunized mice and assessed localization of cells in vivo 4 days post-immunization. In accordance with transcriptomic data, DOT1L-deficiency resulted in altered positioning of B cells within the spleen at the critical time juncture of nascent GC formation in vivo. In *Cd23*^cre/+^ mice, B cell lymphoma-6 (BCL6)^+^ B and T cells were observed in the follicle in nascent GC (Figure 4C; orange arrows), consistent with previous studies (Kerfoot et al., 2011; Kitano et al., 2011). In DOT1L-deficient mice, BCL6^+^ cells were detectable, confirming that upregulation of BCL6 in the early stages of the response could occur in *Dot1l*^*f/f*^*Cd23*^Cre/+^ mice (Figure 4C). Yet in contrast to *Cd23*^cre/+^ mice, DOT1L-deficient BCL6^+^ cells did not cluster together in the follicle. Instead, BCL6^+^ cells in *Dot1l*^*f/f*^*Cd23*^Cre/+^ mice were observed to be located mainly outside the B220^+^ follicles (Figure 4C; white arrows). Specifically, BCL6^+^ cells were located in the outer edge of the follicle, potentially in the marginal zone, as well as in extrafollicular areas (Figures 4C), with quantitation revealing a >3-fold increase in BCL6^+^ cells positioned in the outer edge of the follicle in *Dot1l*^*f/f*^*Cd23*^Cre/+^ mice compared to *Cd23*^cre/+^ mice (Figure 4D). Additionally, more B220^+^ cells were observed to penetrate into the T cell zone in the absence of DOT1L (Figure 4C).

**Figure 4:**
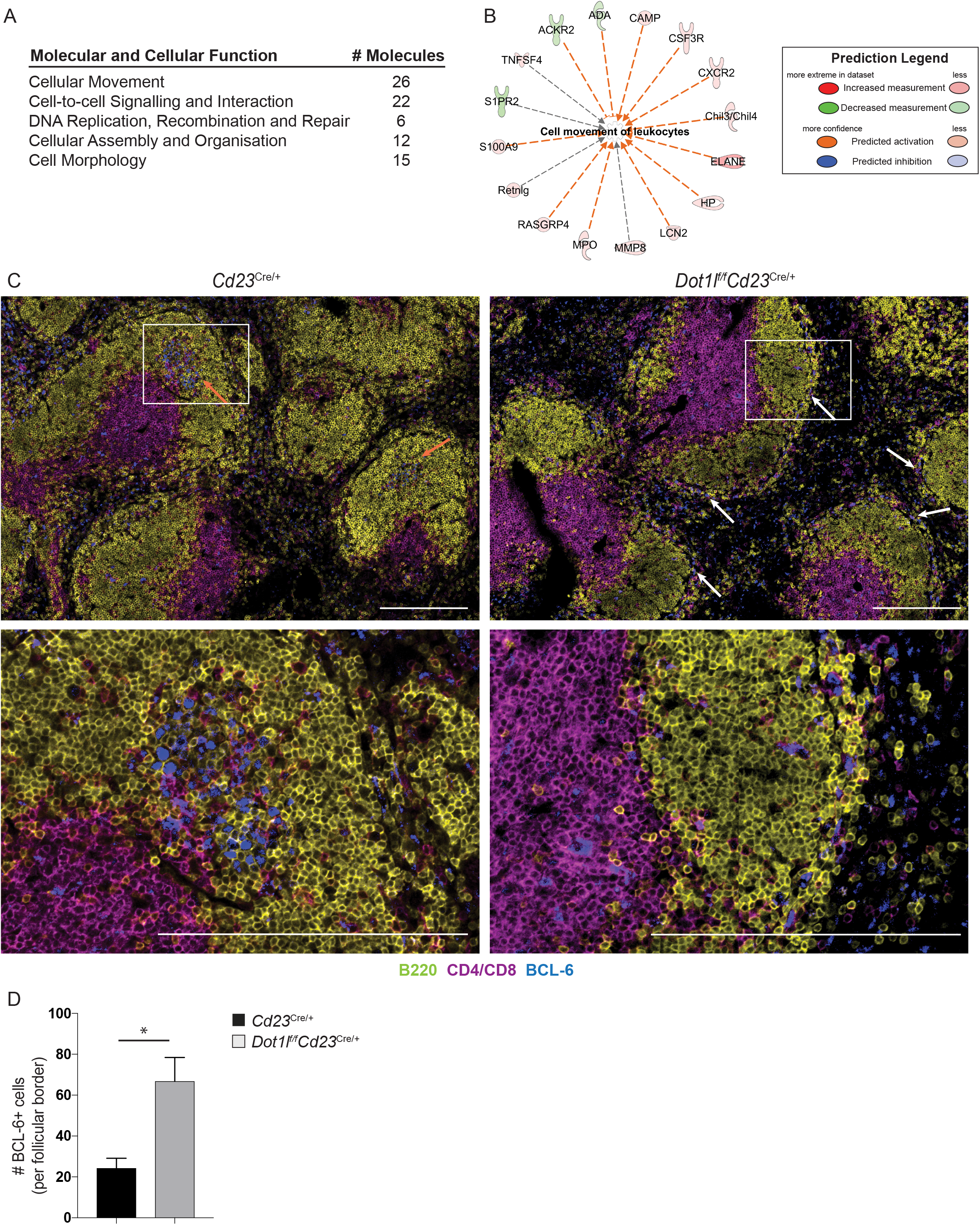
DOT1L regulates localization of activated B cells during a humoral response. (A) IPA was used for the biological functions and pathway analysis of differentially expressed genes in CD19^+^IgD^−^ samples. Summary of top molecular and cellular function regulated by Dot1L. Prediction analysis of potential diseases and functions from sets of identified differentially expressed genes in CD19^+^IgD^−^ samples. Annotation of altered genes involved in cell movement of leukocytes. The different colours indicate the expression level of the genes (red indicating up-regulated genes and green indicating down-regulated genes) or the predicted activity of the identified gene in the activation (blue) or inhibition (orange) of the specified function. The dashed arrows indicate the indirect relationship of the annotated gene to the altered function. (C) Representative images from histological analyses day 4 post-immunization: B220 (yellow), CD4/CD8 (magenta) and BCL6 (blue). Scale bar = 100μm. (D) Quantitation of BCL6^+^ cells in the outer edge of follicles. * P < 0.05.

It was possible that background responses to environmental antigens were the cause of histological differences observed between *Cd23*^cre/+^ and *Dot1l*^*f/f*^*Cd23*^Cre/+^ mice. Therefore, we subcutaneously immunised *Cd23*^cre/+^ and *Dot1l*^*f/f*^*Cd23*^Cre/+^ mice with NP-KLH in alum and assessed draining and non-draining lymph nodes to confirm the location of B cells responding to the immunogen. At day 5 post-immunization, no BCL6^+^ cells were detectable in the non-draining lymph node (data not shown). In contrast, nascent GCs were detected in *Cd23*^cre/+^ mice in the draining lymph node, whereas no clusters of BCL6^+^ cells in *Dot1l*^*f/f*^*Cd23*^Cre/+^ mice were detected in the follicle, but instead at the outer edge of the follicle (Figure S4H), concordant with the splenic results. Thus, dysregulation of a network of genes involved in cellular migration in *Dot1l*^*f/f*^*Cd23*^Cre/+^ mice correlated with altered localization in the spleen and lymph node during the initial stages of an immune response. Taken together, this study establishes DOT1L as a pivotal regulator of B cell development, antigen-driven B cell migration, differentiation and the establishment of long-lived humoral immune memory.

Elucidation of the critical histone modifiers that regulate B cell fate and function has significant implications for translational studies. In particular, dissecting the role of DOT1L in cellular biology is vital for understanding the biological effects of clinical intervention with DOT1L inhibitors. A DOT1L small molecule inhibitor is currently in clinical trials for the treatment of *MLLr* leukemias (Stein and Tallman, 2015). Our results suggest that sustained inhibition of DOT1L could have an impact on B cell progenitors and the ability of the humoral system to respond to infection. Further, the critical role of DOT1L in multiple stages of B cell biology may indicate that DOT1L could be an important target for other B cell-derived cancers. For example, DOT1L was identified in a screen of patients with diffuse large B cell lymphoma (Szablewski et al., 2018). This raises the possibility that targeting DOT1L in GC B cell-derived lymphomas may be a potential, but as yet unexplored, novel therapeutic option.

In summary, we have identified DOT1L as a critical and central regulator of B cell biology. Here, we demonstrate for the first time that appropriate B cell development, as well as migration and ultimately differentiation during immune responses, requires DOT1L function. This work opens up new avenues of investigation in both understanding the fundamental molecular underpinnings of B cell responses and whether small molecule inhibitors targeting DOT1L may be effective in abrogating B cell-driven diseases.

## Supporting information

Key Resources Table

## Abbreviations used

ASCs: antibody-secreting cells
BCL6: B cell lymphoma-6
CPM: counts per million
CTV: cell trace violet
DOT1L: disruptor of telomeric silencing 1-like
EZH2: enhancer of zeste homolog 2
FVS: fixable viability stain
H3K27me3: trimethylation of lysine 27 on histone H3
Ig: immunoglobulin
KLH: Keyhole Limpet Hemocyanin
MLLr-ALL: Mixed Lineage Leukaemia-rearranged acute lymphoblastic leukaemia
NP: (4-Hydroxy-3-nitrophenyl)-acetyl
PRC2: polycomb repressive complex 2
S1PR: sphingosine-1-phosphate receptor
Th cells: T helper cells

## ACKNOWLEDGEMENTS

We thank Mireille Lahoud and David Tarlinton for critical reading of this manuscript; members of the Good-Jacobson and Zaph labs, Monash Micromon and Bioinformatics Platforms for technical assistance. KLG-J is a Bellberry-Viertel Senior Medical Research Fellow and was supported by a National Health and Medical Research Council (NHMRC) Career Development Fellowship. ADiP was supported by an American Association of Immunologists Careers in Immunology Fellowship. This work was supported by a NHMRC project grant to JRG and KLG-J (GNT1137989). JRG was supported by a Walter and Eliza Hall Institute Centenary Fellowship sponsored by CSL. LK was supported by a Monash University Research Training Program Scholarship. LD was supported by a University of Melbourne research scholarship. CZ was funded by NHMRC Project grants GNT1104433 and GNT1104466. CZ is a veski Innovation Fellow.

## AUTHOR CONTRIBUTIONS

KLG-J designed the research; LK, ADiP, LH and LD performed research; JRG performed research and provided intellectual input; SS and CZ provided reagents, mice and intellectual input; and KLG-J wrote the manuscript.

## STAR METHODS

### 1. LEAD CONTACT AND MATERIALS AVAILABILITY

This study did not generate new unique reagents.

Further information and requests for resources and reagents should be directed to and will be fulfilled by the Lead Contact, Kim Good-Jacobson.

### 2. EXPERIMENTAL MODEL AND SUBJECT DETAILS

#### Mice, immunizations and purification of cells

*Mb1-Cre* (Pelanda et al., 2002), *Eed^f/f^* (Xie et al., 2014) and *Cd23*-Cre (Kwon et al., 2008) were provided by Michael Reth, Stuart Orkin and Meinrad Busslinger, respectively. To create *Dot1l^f/f^* mice, we derived mice from DOT1L targeted ES cells (Dot1ltm1a(KOMP)Wtsi) from UCDavis KOMP Repository and crossed them with FLP mice (Monash University). Subsequently, *Dot1l^f/f^* mice were crossed with *Mb1-Cre* or *Cd23-Cre* (all on C57BL/6 background). Animal procedures were approved by Monash University Animal Ethics Committee and all mice were maintained at the Monash Animal Research Platform. Mice at least 6 weeks of age and of either gender were used in this study. Mice were humanely euthanized by hypercapnea. *T-dependent immunization model:* (4-Hydroxy-3-nitrophenyl)-acetyl (NP) hapten was conjugated to Keyhole Limpet Hemocyanin (NP_13_KLH), precipitated on 10% Alum and diluted in sterile PBS to a final concentration of 100μg/100μl. *T-independent immunization model:* NP_55_Ficoll was diluted in sterile PBS to a final concentration of 40μg/100μl. *Influenza infections:* Mice were inoculated with 250p.f.u. of HKx31 (H3N2) influenza virus, generously provided by Gabrielle Belz and Stephen Turner, as previously described (Belz et al., 2000; Flynn et al., 1998). *Anasthesia*: Isofluorane (2.5%) was used to lightly anesthetize mice for intranasal infections. Mice were monitored to ensure stabilization of breathing and were warmed by a lamp to alleviate suffering if necessary.

### 3. METHOD DETAILS

#### Flow cytometry

Single cells were resuspended in PBS 2% FCS and stained for flow cytometric analysis as previously described (Cooper et al., 2018). Briefly, 5 × 10^6^ cells were resuspended in 50μl of FVS fixable viability stain, diluted 1:1000 in PBS 2% FCS and incubated for 15 minutes at room temperature in the dark. Samples were then washed with PBS 2% FCS, and then resuspended in 50μl of indicated antibodies (Table S3) and incubated at 4°C for 20 minutes. FcγRII/III (24G2; supernatant) and normal rat serum (Sigma-Aldrich) was used to block non-specific binding. Cells were then washed and resuspended in PBS 2% FCS and data acquired on a BD Fortessa or LSRIIa and subsequently analyzed using FlowJo (Tree Star). *Sort-purification*: cells were stained with antibodies and purified by FACS Influx (BD), with purity >98%.

#### ELISPOT

ASCs of the IgG1, IgG3 and IgM isotype were analyzed by ELISPOT. Multiscreen HA plates (Millipore) were coated overnight at 4°C with NP_12_BSA for the quantification of NP-specific IgG1^+^ and IgG3^+^ ASCs. An NP_9_BSA coat was used to analyze total NP^+^IgM^+^ ASCs while NP_1_BSA was used to measure high-affinity IgG1^+^ ASCs. Plates were blocked with PBS 1%BSA for 1 hour, washed with PBS, and loaded with samples prepared in RPMI 5%FCS, 50μM 2-Mercaptoethanol and 2mM Glutamine before an overnight incubation at 37°C. Plates were washed with PBS-Tween and distilled water and subsequently incubated for 1 hour at 37°C with secondary antibody (IgG1, IgG3 or IgM) conjugated to alkaline-phosphatase (Southern Biotech) for one hour. Plates were washed, as before, and developed with the BCiP®/NBT reaction (Sigma-Aldrich).

#### ELISA

Serum samples from mice immunized with NP-KLH or NP-Ficoll were assessed by coating 96-well high binding plates (Sarstedt), overnight at 4°C, with 5μg/μl NP_9_BSA for IgM^+^ antibodies or NP_12_BSA for all other isotypes. Serum samples from naïve mice or influenza-infected mice were analyzed by coating plates with the appropriate isotype-specific capture antibody (Southern Biotech; Table S3). Plates were blocked, the following day, with PBS 1%BSA for 1 hour, washed with PBS-Tween and distilled water and then loaded with serially-diluted serum samples before a 4-hour incubation at 37°C. Plates were washed again with PBS-Tween and distilled water and then incubated for 1 hour at 37°C with the appropriate anti-mouse secondary antibody conjugated to horseradish peroxidase (Southern Biotech; Table S3). Plates were washed and developed with OPD-substrate solution (Sigma-Aldrich). Serum was serially diluted in block and the optical density values for each sample are shown as a non-linear regression (sigmoidal curve fit).

#### Immunohistochemistry

Spleens from mice immunized with NP-KLH were extracted, 7 days post-immunization, and frozen in OCT (Tissue-Tek). 7μm sections were cut using a microtome (Leica) at −20°C and fixed with acetone. Staining was performed using described antibodies (Table S3). Slides were viewed under 4X and 10X magnification using an Olympus CKX41 microscope and images were captured with a mounted Nikon DP-12 camera. Raw images were taken using the Nikon NIS-Element software platform while the processing program, ImageJ, was used to add 100μm scale bars to each image.

#### Immunofluorescence

Spleens or inguinal lymph nodes were either directly embedded in OCT compound (Tissue-Tek), or first fixed in 4% paraformaldehyde and immersed in 30% sucrose before being embedded in OCT compound (Tissue-Tek). Tissues were cut via microtome (Leica) into 12μm sections and mounted on Superfrost Plus slides. Sections were fixed with cold acetone (Sigma) and stained as previously described (Rankin et al., 2013) and using indicated antibodies (Table S3). Images were acquired using a LSM780 confocal microscope (Carl Zeiss MicroImaging). The acquisition software was Zen Black 2012.

#### Cell Culture

*Deletion model in culture:* CD19^+^ naïve B cells were isolated using a B cell negative enrichment kit (STEMCELL) from *Cd23*^Cre/+^ mice and *Dot1l*^*f/f*^*Cd23*^Cre/+^ mice. B cells were labeled in PBS with 2 μM CTV (Life Technologies) for 10 minutes at 37°C. Cells were then washed and resuspended in RPMI 5%FCS, 50μM 2-Mercaptoethanol and 2mM Glutamine. 5 × 10^4^ labelled B cells were cultured with 50ng/mL CD40L (R&D Systems) in combination with 50ng/mL IL-4 and 5ng/mL IL-5 (R&D Systems). *DOT1L inhibitor in culture:* B cells were purified from wildtype mice as above. Purified B cells were labelled with CTV and cultured with LPS and IL-4. Specified wells were treated with either DMSO or a DOT1L inhibitor (SGC0946) diluted in media.

#### Chemotaxis assay

B cells were isolated using negative B cell isolation kit (Stem Cell) and resuspended in 0.5% FBS, RPMI 1640 at 5×10^5^cells per well. 100μl/well were added to the upper chamber of transwell plates and transmigrated across 3μm transwell filters (Corning Costar Corp, NY, USA) for 3 hours. Chemotaxis agent CCL21 (Peprotech, NJ, USA) diluted in 0.5% FBS, RPMI 1640 was added to the bottom chamber at indicated concentrations. Post migration, cells were collected, stained for surface markers (Table S3) and analysed and enumerated by flow cytometry.

#### RNA-sequencing and bioinformatics analysis

Sort-purified cell subsets were centrifuged, lysed in RLP Buffer (QIAGEN) and passed through a gDNA eliminator spin column (QIAGEN) to remove genomic DNA. The flow-through was passed through a silica-membrane designed to capture RNA (QIAGEN), which was then washed and dried following the manufacturer’s recommendations (QIAGEN RNeasy Plus Micro Kit), and eluted in 14μl of RNase-free H2O. The 8 RNA samples were prepped using the Illumina Truseq stranded mRNA sample preparation kit with the input RNA of 100ng and sequence using Illumina Nextseq 500 Single End 75 cycles high output version2 cartridge. Raw fastq files were analyzed using the RNAsik pipeline (Tsyganov et al., 2018) using STAR aligner (Dobin et al., 2013) with the GRCm38.p4 Ensembl reference. Reads were quantified using featureCounts (Liao et al., 2014) producing the raw genes count matrix and various quality control metrics. Raw counts were then analyzed with Degust (Powell, 2015), a web tool which performs differential expression analysis using limma voom normalisation (Law et al., 2014), producing counts per million (CPM) library size normalisation and trimmed mean of M values normalisation (Robinson and Oshlack, 2010) for RNA composition normalisation. Degust (Powell, 2015) also provides quality plots including classical multidimensional scaling and MA plots. Data is shown with only protein coding genes and with immunoglobulin genes removed. IPA (QIAGEN) was used for the biological functions and pathway analysis of differentially expressed genes.

#### Histone Extractions and Western Blotting

1×10^6^ cells were isolated per sample using fluorescence-activated cell sorting and lysed, overnight at 4°C, in 0.2M HCl. Cell debris was pelleted and the recovered supernatant was treated at 95°C, for 5 min, in SDS-PAGE loading buffer (250mM Tris-HCl pH 6.8, 10% SDS, 50% Glycerol, 0.05% bromophenol blue, 5% 2-mercaptoethanol). Samples were resolved on a 12% SDS-PAGE and analyzed by western blot using anti-H3 (Abcam ab1791), anti-H3K27me3 (Abcam ab6002) and anti-H3K79me2 (Abcam ab3594) antibodies. Blots were further stained with Horseradish Peroxidase- conjugated goat anti-mouse (Southern Biotech) or anti-rabbit IgG (H + L) antibodies (Biorad), developed with Clarity Western ECL Substrate (Biorad) and scanned on the ChemiDoc™ Touch Imaging System (Biorad).

### 4. QUANTIFICATION AND STATISTICAL ANALYSIS

The Mann-Whitney nonparametric, two-tailed test was used for statistical analyses of all data, with the exception of western blot data, using GraphPad Prism software. Paired t-test was used for statistical analyses of western blot data, using GraphPad Prism software. All data is presented as the mean +/− the standard error of the mean. n = the number of individual mice assessed.

### 5. DATA AND CODE AVAILABILITY

The accession number for the RNA-seq data reported in this paper is GEO: https://www.ncbi.nlm.nih.gov/geo/query/acc.cgi?acc=GSE138401

### 6. KEY RESOURCES TABLE

In separate attachment.

## SUPPLEMENTAL INFORMATION

***Figure S1: Conditional Dot1l deletion disrupts B cell development. Related to Figure 1.***

(A) Schematic of conditional deletion of *Dot1l* in the B cell lineage. (B) Representative image of spleens isolated from *Mb1^Cre/+^* (left) and *Dot1l^f/f^ Mb1^Cre/+^* (right) mice. (C) Spleen weight of naïve, adult *Mb1^Cre/+^* (black bars) and *Dot1l^f/f^Mb1^Cre/+^* mice (light grey bars). Results are combined from 2 independent experiments. (D) Flow cytometric analyses of CD19^+^ B cells in the spleen of naïve, adult *Mb1^Cre/+^* (black bars), *Dot1l^fl+^ Mb1^Cre/+^* (grey bars) and *Dot1l^f/f^Mb1^Cre/+^* mice (light grey bars). (E) Representative images from histological analyses of spleens from naïve mice: MOMA-1 (yellow) and B220 (magenta). Scale bar = 100μm. (F-I) Frequency and number of (F-G) marginal zone B cells and (H-I) follicular B cells in the spleen. (J-O) Different antibody isotypes from the serum of *Mb1^Cre/+^* (black squares) and *Dot1l^f/+^* and *Dot1l^f/f^ Mb1^Cre/+^* mice (white circles) were compared by ELISA analysis: (J) total IgM, (K) total IgG1, (L) total IgG2c, (M) total IgG2b, (N) total IgG3 and (O) total IgA serum levels for each mouse group (n=6). Sigmoidal curve fit of serially diluted samples. (P-Q) Flow cytometric analyses of B cell populations in the bone marrow of naïve, adult *Mb1^Cre/+^* (black bars), *Dot1l^fl+^ Mb1^Cre/+^* (grey bars) and *Dot1l^f/f^Mb1^Cre/+^* mice (light grey bars). (P) B220^+^ populations, (Q) Pro B cell (B220^lo^CD24^+^BP1^−^CD43^+^IgM^−^), Pre B cell (B220^lo^CD24^+^BP1^+^CD43^−^IgM^−^IgD^−^), Immature B cell (B220^+^CD24^+^CD43^−^IgM^lo^IgD^−^), Transitional B cell (B220^+^CD24^+^CD43^−^IgM^hi^IgD^−^), Early Mature B cell (B220^+^CD24^+^CD43^−^IgM^hi^IgD^+^) and Late Mature B cell (B220^+^CD24^+^CD43^−^IgM^lo^IgD^+^) populations in the bone marrow. *Mb1^Cre/+^* (n=8), *Dot1l^f/+^Mb1^Cre/+^* (n=5) and *Dot1l^f/f^Mb1^Cre/+^* (n=7) mice. Results are combined from four independent experiments. * P < 0.05; ** P < 0.01; *** P < 0.001.

***Figure S2: Regulation of B cells in the absence of DOT1L. Related to Figures 1 and 2.***

(A) Flow cytometric analyses of follicular B cell frequency in naïve *Cd23*^Cre/+^ and *Dot1l*^*f/f*^*Cd23*^Cre/+^ mice. (B) Flow cytometric analyses of marginal zone B cells. n=5 per group. (C) *Cd23*^Cre/+^ and *Dot1l*^*f/f*^*Cd23*^Cre/+^ mice were intranasally infected with HKx31 and culled for analysis 8 days and 14 days post-infection. Flow cytometric representative plot of CD95^+^CD38^lo^ splenic GC B cells. (D-E) Frequency (D) and (E) number of CD95^+^CD38^lo^ GC B cells in the spleen. (F-G) Frequency (F) and (G) number of CD95^+^CD38^lo^ GC B cells in the mediastinal lymph node. (H) Frequency of splenic and mediastinal lymph nodes GC B cells that have switched to IgG2c. n=6 per genotype, combined from two independent experiments per time point. * P < 0.05; ** P < 0.01. (I) ELISA analysis of IgG1 serum antibody from *Cd23^Cre/+^* (closed squares) and *Dot1l^f/f^ Cd23^Cre/+^* mice (open circles) infected with influenza. (J) ELISA analysis of NP^+^IgG2c serum antibody from *Cd23^Cre/+^* and *Dot1l^f/f^ Cd23^Cre/+^* mice immunized with NP-KLH in alum. ** P < 0.01. (K) CD19^+^ B cells from either *Dot1l*^*f/f*^*Cd23*^Cre/+^ or *Cd23*^Cre/+^ mice were CTV-labelled stimulated in vitro with CD40L, IL-4 and IL-5 for 3.5d and assessed by flow cytometry. Frequency of cells in each division, determined by CTV division peaks, is shown. (L) CD19^+^ B cells purified from the spleens of C57Bl/6 mice were labelled with CTV, stimulated with LPS and IL-4 incubated with small molecule inhibitor SGC946 to DOT1L, compared to a DMSO control. Frequency of cells per division is shown.

***Figure S3: Regulation of global histone modifications in B cells by DOT1L and PRC2. Related to Figures 1 and 2.***

(A) Schematic of assessment of H3K79me2 and H3K27me3: mice were immunized with NP-KLH in alum and CD19^+^IgD^+^ and CD19^+^IgD^−^ B cell subsets sort-purified post-immunization. (B) Assessment of global H3K79me2 in sort-purified B cell subsets. (C) Quantitation of global H3K79me2. (D) Assessment of global H3K27me3 in sort-purified B cell subsets. (E) Quantitation of global H3K27me3. Blot images are cropped to show area around either H3K27me3 or H3K79me2 bands. No lanes were removed. n=3 per group, combined from three independent experiments. * P < 0.05; ** P < 0.01.

***Figure S4: Modulation of genes in the absence of DOT1L in mature B cell subsets. Related to Figures 3 and 4.***

(A) Multidimensional scaling plot of each sample assessed by RNA-sequencing. (B) Venn Diagram of differential expressed genes was obtained using the Comparison Analysis tool in IPA software to compare different sets of naive and activated B cells. (C-E) Bar graphs of genes differentially expressed solely in the CD19^+^IgD^+^ B cell population (C), common differential genes modulated in both B cell subsets (D), those solely in the CD19^+^IgD^−^ B cell population (E). (F) Migration of CD19^+^IgD^−^NP^+^ splenic B cells towards CCL21. B cells were isolated from either *Dot1l*^*f/f*^*Cd23*^Cre/+^ or *Cd23*^Cre/+^ mice 4 days post-immunization with NP-KLH in alum. Error bars are mean +/-the standard error of the mean. (G) Prediction analysis of potential diseases and functions from sets of identified differentially expressed genes in CD19^+^IgD^−^ samples. Annotation of altered genes involved in the regulation of B cells numbers. The different colours indicate the expression level of the genes (red indicating up-regulated genes and green indicating down-regulated genes) or the predicted activity of the identified gene in the activation (blue) or inhibition (orange) of the specified function. The dashed arrows indicate the indirect relationship of the annotated gene to the altered function. (H) *Dot1l*^*f/f*^*Cd23*^Cre/+^ or *Cd23*^Cre/+^ mice subcutaneously immunized with NP-KLH in alum. Representative images from histological analyses of inguinal lymph nodes 5 days post-immunization: B220 (yellow), CD4/CD8 (magenta) and BCL6 (blue). Scale bar = 100μm.

***Table S1: Differential gene expression in CD19^+^IgD^+^ B cells. Related to Figure 3.***

***Table S2: Differential gene expression in CD19^+^IgD^−^ B cells. Related to Figure 3.***

