## Supplementary material for "The histone methyltransferase DOT1L is essential for humoral immune responses": Key Resources Table

| REAGENT or RESOURCE | SOURCE | IDENTIFIER |
|---|---|---|
| Antibodies | | |
| Anti-CD138 PE (clone 281-2) | BioLegend | Cat # 142504; RRID:AB_10916119 |
| Anti-CD138 BV605 (clone 281-2) | BD Bioscience | Cat # 563147; RRID:AB_2721029 |
| Anti-CD45R (B220) AF488 (clone RA3-6B2) | BioLegend | Cat # 103225; RRID:AB_389308 |
| Anti-CD45R (B220) APC-Cy7 (clone RA3-6B2) | BioLegend | Cat # 103224; RRID:AB_313007 |
| Anti-CD45R (B220) AF555 (clone RA3-6B2) | Produced in-house | N/A |
| Anti-IgG1 APC (clone X56) | BD Bioscience | Cat # 550874; RRID:AB_398470 |
| Anti-CD19 APC-Cy7 (clone 6D5) | BioLegend | Cat #115530; RRID:AB_830707 |
| Anti-CD19 Pacific Blue (clone 6D5) | BioLegend | Cat # 115523; RRID:AB_439718 |
| Anti-CD19 PE (clone 6D5) | BioLegend | Cat # 115508; RRID:AB_313643 |
| Anti-IgD AF488 (clone 11-26c.2a) | BioLegend | Cat # 405718; RRID:AB_10730619 |
| Anti-IgD PerCP-Cy5.5 (clone 11-26c.2a) | BD Bioscience | Cat # 564273; RRID:AB_2738722 |
| Anti-CD95 PE-Cy7 (clone Jo2) | BD Bioscience | Cat # 557653; RRID:AB_396768 |
| Anti-CD38 Pacific Blue (clone 90) | BioLegend | Cat # 102720; RRID:AB_10613468 |
| Anti-IgG2a/2b FITC (clone R2-40) | BD Bioscience | Cat # 553399; RRID:AB_394837 |
| Anti-CD43 APC (clone S11) | BioLegend | Cat # 143207; RRID:AB_11149489 |
| Anti-CD249 (BP1) PE (clone BP-1) | BD Bioscience | Cat # 553735; RRID:AB_395018 |
| Anti-CD24 BV421(clone M1/69) | BioLegend | Cat # 101825; RRID:AB_10901159 |
| Anti-IgM AF488 (clone RMM-1) | BioLegend | Cat # 406522; RRID:AB_2562859 |
| Anti-CD169 (MOMA-1) FITC (clone 3D6.112) | Bio-Rad | Cat # MCA884F; RRID:AB_324246 |
| Anti-CD23 AF488 (clone B3B4) | BioLegend | Cat # 101609; RRID:AB_493362 |
| Anti-CD21/CD35 APC (clone 7G6) | BioLegend | Cat # 123412; RRID:AB_2085160 |
| Anti-CD11b APC-Cy7 (clone M1/70) | TONBO | Cat # 25-0112; RRID:AB_2621625 |
| Anti-CD3e PE-Cy7 (clone 145-2C11) | eBioscience | Cat # 25-0031-82; RRID:AB_469572 |
| Anti-CD4 A488 (clone GK1.5-7) | Produced in-house | N/A |
| Anti-CD8a A488 (clone 53.6.7) | Produced in-house | N/A |

| | | |
|---|---|---|
| Anti-Bcl6 A647 (clone K112-91) | BD Pharmingen | Cat #561525; RRID:AB_10898007 |
| Goat anti-mouse IgG1 unlabeled (polyclonal) | Southern Biotech | Cat # 1070-01; RRID:AB_2794408 |
| Goat anti-mouse IgG2a unlabeled (polyclonal) | Southern Biotech | Cat # 1080-01; RRID:AB_2794475 |
| Goat anti-mouse IgG2b unlabeled (polyclonal) | Southern Biotech | Cat # 1090-01; RRID:AB_2794517 |
| Goat anti-mouse IgG3 unlabeled (polyclonal) | Southern Biotech | Cat # 1100-01; RRID:AB_2794567 |
| Goat anti-mouse IgA unlabeled (polyclonal) | Southern Biotech | Cat # 1040-01; RRID:AB_2314669 |
| Goat anti-mouse IgM unlabeled (polyclonal) | Southern Biotech | Cat # 1020-01; RRID:AB_2794197 |
| Goat anti-mouse IgG1 HRP (polyclonal) | Southern Biotech | Cat # 1070-05; RRID:AB_2650509 |
| Goat anti-mouse IgG2a HRP (polyclonal) | Southern Biotech | Cat # 1080-05; RRID:AB_2734756 |
| Goat anti-mouse IgG2c HRP (polyclonal) | Southern Biotech | Cat # 1079-05; RRID:AB_2794466 |
| Goat anti-mouse IgG2b HRP (polyclonal) | Southern Biotech | Cat # 1090-05; RRID:AB_2794521 |
| Goat anti-mouse IgG3 HRP (polyclonal) | Southern Biotech | Cat # 1100-05; RRID:AB_2794573 |
| Goat anti-mouse IgA HRP (polyclonal) | Southern Biotech | Cat # 1040-05; RRID:AB_2714213 |
| Goat anti-mouse IgM HRP (polyclonal) | Southern Biotech | Cat # 1020-05; RRID:AB_2794201 |
| Goat anti-mouse IgG3 AP (polyclonal) | Southern Biotech | Cat # 1100-04; RRID:AB_2794572 |
| Goat anti-mouse IgG1 AP (polyclonal) | Southern Biotech | Cat # 1070-04; RRID:AB_2794411 |
| Goat anti-mouse IgM AP (polyclonal) | Southern Biotech | Cat # 1020-04; RRID:AB_2794200 |
| Rat anti-mouse CD45R (B220) (clone RA3-6B2) | BD Pharmingen | Cat # 550286; RRID:AB_393581 |
| AffiniPure Goat Anti-Rat IgG (H+L) (polyclonal) | Jackson Laboratory | Cat # 112035003; RRID:AB_2338128 |
| Streptavidin Alkaline Phosphatase | Southern Biotech | Cat # 7100-04 |
| Anti-H3 (polyclonal) | Abcam | Cat# ab1791; RRID:AB_302613 |
| Anti-H3K27me3 (clone 6002) | Abcam | Cat# ab6002; RRID:AB_305237 |
| Anti-H3K79me2 (polyclonal) | Abcam | Cat# ab3594; RRID:AB_303937 |
| AffiniPure Goat Anti-Rabbit IgG (H+L) HRP (polyclonal) | Southern Biotech | Cat # 4050-05; RRID:AB_2795955 |
| AffiniPure Goat Anti-Mouse IgG (H+L) HRP (polyclonal) | Biorad | Cat # 1706516; RRID:AB_11125547 |
| Bacterial and Virus Strains | | |
| Influenza virus strain HKx31 (H3N2) | Produced in house | N/A |
| Biological Samples | | |

| | | |
|---|---|---|
| Mouse Tissue | N/A | N/A |
| **Chemicals, Peptides, and Recombinant Proteins** | | |
| NP-PE | Conjugated in-house | N/A |
| Peanut Agglutinin | Vector Laboratories | Cat # B-1075 |
| CD40L | R&D Systems | Cat # 8230-CL-050 |
| IL4 | R&D Systems | Cat # 405-ML-010 |
| IL5 | R&D Systems | Cat # 405-ML-005 |
| Cell Trace Violet | Invitrogen | Cat # C34557 |
| SGC0946 | SGC | N/A |
| BD Cytofix™ | BD Biosciences | Cat#554714 |
| Clarity Max™ Western ECL Substrate | Biorad | Cat # 1705062; RRID:AB_2036911 |
| O.C.T Compound | Tissue-Tek | Cat# 4583 |
| **Critical Commercial Assays** | | |
| Fixable Viability Stain 700 | BD Horizon | Cat # 564997 |
| AEC Peroxidase (HRP) Substrate Kit | Vector Laboratories | Cat # SK 4200 |
| Vector® Blue Alkaline Phosphatase Substrate Kit | Vector Laboratories | Cat # SK 5300 |
| **Deposited Data** | | |
| RNA-sequencing data | This paper | https://www.ncbi.nlm.nih.gov/geo/query/acc.cgi?acc=GSE138401 |
| **Experimental Models: Organisms/Strains** | | |
| *Mb1-Cre* | Pelanda et al., 2002 | N/A |
| *Eed*$^{f/f}$ | Xie et al., 2014 | N/A |
| *Cd23*-Cre | Kwon et al., 2008 | N/A |
| Dot1ltm1a(KOMP)Wtsi | UCDavis KOMP Repository | CSD29070 |
| **Software and Algorithms** | | |
| Flowjo (Treestar) | FlowJo, LLC | https://www.flowjo.com/ |
| Ingenuity Pathway Analysis | QIAGEN | N/A |
| Degust | Powell, 2015 | http://degust.erc.monash.edu/ |
| NIS-Elements | Nikon | https://www.nikon.com/products/microscope-solutions/lineup/img_soft/nis-elements/ |
| ImageJ | National Institutes of Health (NIH) | https://imagej.nih.gov/ij/index.html |
| Prism | Graph Pad | https://www.graphpad.com/scientific-software/prism/ |
| Zen Black | ZEISS | https://www.zeiss.com/microscopy/int/products/microscope-software/zen.html |
| **Other** | | |
| Superfrost Plus Adhesion Microscope Slides | Thermo Scientific | Cat# J1800AMNT |

CellPress
