## Supplemental Information for "The histone methyltransferase DOT1L is essential for humoral immune responses"

Figure S1

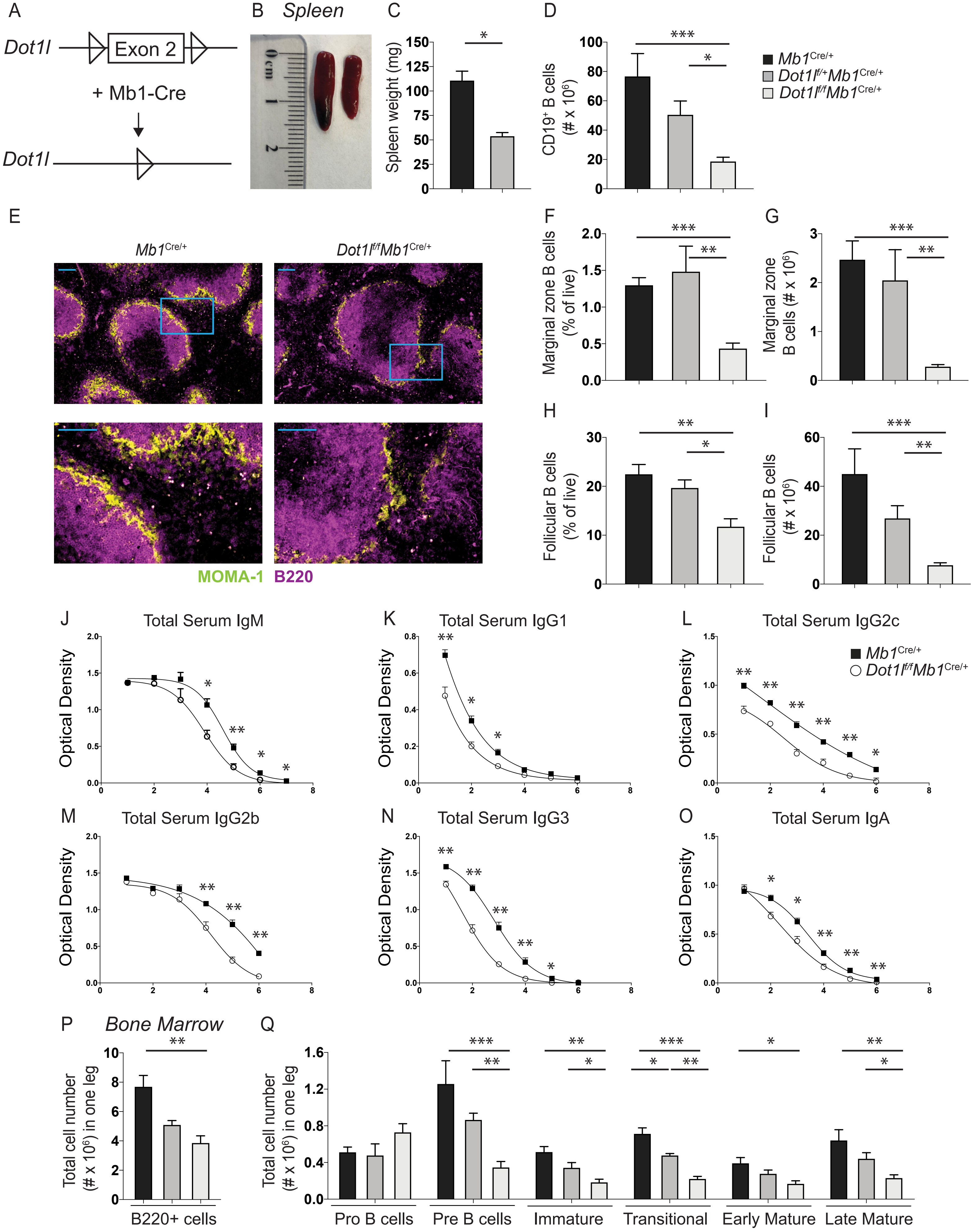
A
Dot1l
Exon 2
+ Mb1-Cre
Dot1l
B Spleen
C
Spleen weight (mg)
D
CD19+ B cells (# x 10^6)
Mb1Cre/+
Dot1lf/+Mb1Cre/+
Dot1lf/fMb1Cre/+
E
Mb1Cre/+
Dot1lf/fMb1Cre/+
MOMA-1 B220
F
Marginal zone B cells (% of live)
G
Marginal zone B cells (# x 10^6)
H
Follicular B cells (% of live)
I
Follicular B cells (# x 10^6)
J Total Serum IgM
K Total Serum IgG1
L Total Serum IgG2c
Mb1Cre/+
Dot1lf/fMb1Cre/+
M Total Serum IgG2b
N Total Serum IgG3
O Total Serum IgA
Optical Density
P Bone Marrow
Total cell number (# x 10^6) in one leg
B220+ cells
Q
Total cell number (# x 10^6) in one leg
Pro B cells
Pre B cells
Immature
Transitional
Early Mature
Late Mature

Figure S2

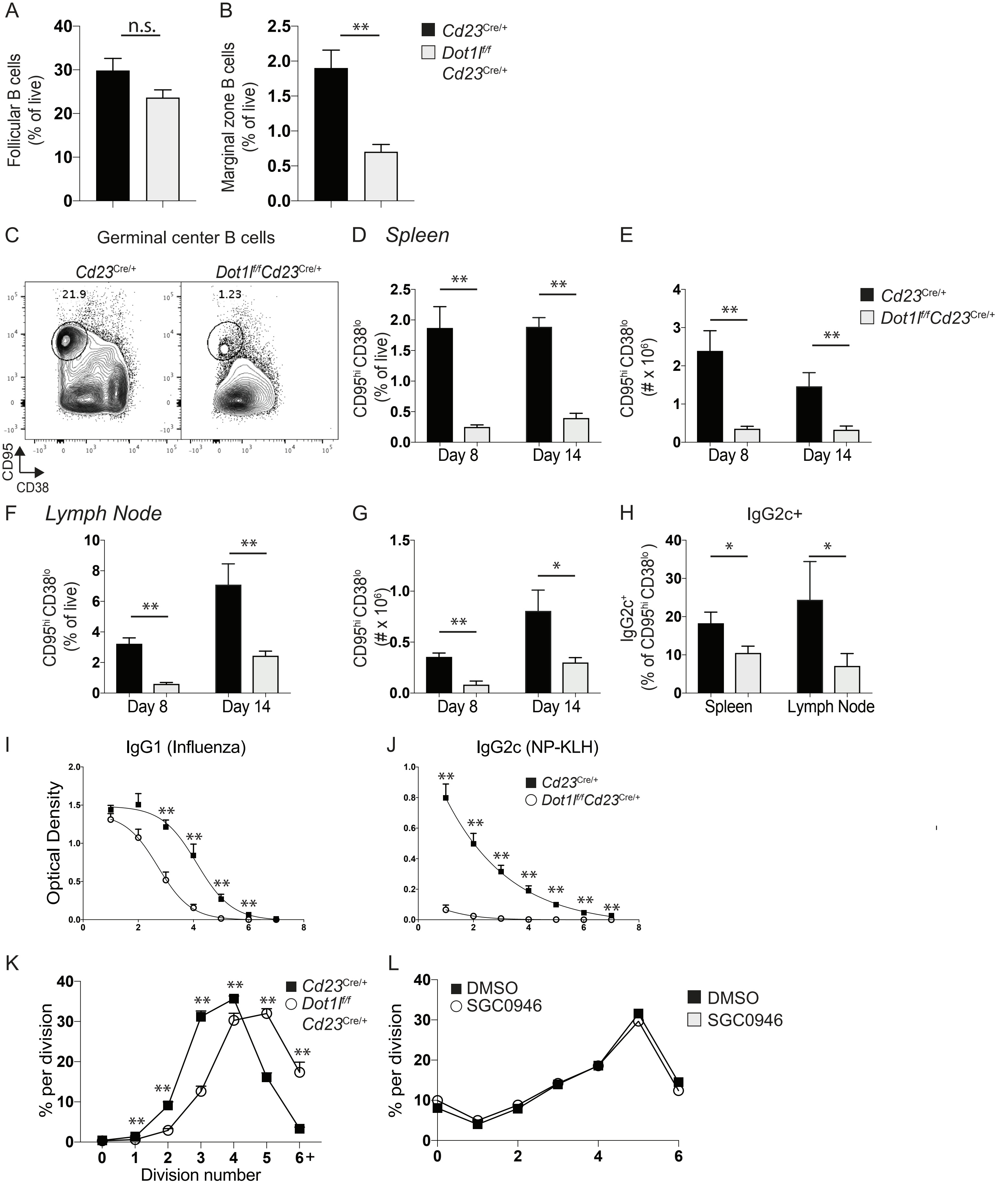

A
B
C
D
E
F
G
H
I
J
K
L
n.s.
Follicular B cells (% of live)
Marginal zone B cells (% of live)
Cd23Cre/+
Dot1lf/f Cd23Cre/+
Germinal center B cells
21.9
1.23
CD95
CD38
Spleen
CD95hi CD38lo (% of live)
CD95hi CD38lo (# x 10^6)
Day 8
Day 14
Lymph Node
IgG2c+
IgG2c+ (% of CD95hi CD38lo)
Spleen
Lymph Node
IgG1 (Influenza)
IgG2c (NP-KLH)
Optical Density
% per division
Division number
DMSO
SGC0946

Figure S3

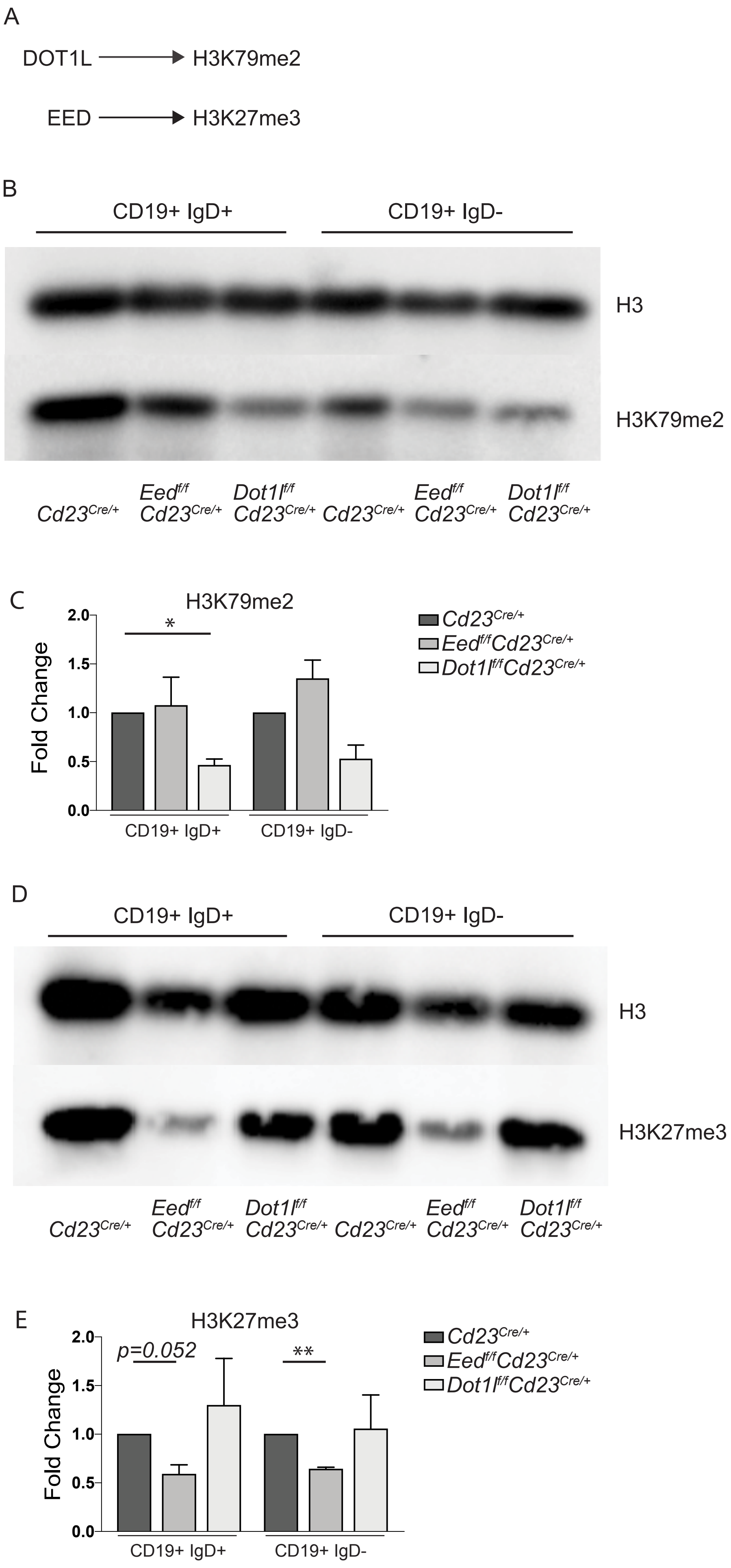
A
DOT1L
H3K79me2
EED
H3K27me3
B
CD19+ IgD+
CD19+ IgD-
H3
H3K79me2
Cd23Cre/+
Eedf/f Cd23Cre/+
Dot1lf/f Cd23Cre/+
Cd23Cre/+
Eedf/f Cd23Cre/+
Dot1lf/f Cd23Cre/+
C
H3K79me2
Fold Change
*
CD19+ IgD+
CD19+ IgD-
Cd23Cre/+
Eedf/fCd23Cre/+
Dot1lf/fCd23Cre/+
D
CD19+ IgD+
CD19+ IgD-
H3
H3K27me3
Cd23Cre/+
Eedf/f Cd23Cre/+
Dot1lf/f Cd23Cre/+
Cd23Cre/+
Eedf/f Cd23Cre/+
Dot1lf/f Cd23Cre/+
E
H3K27me3
Fold Change
p=0.052
**
CD19+ IgD+
CD19+ IgD-
Cd23Cre/+
Eedf/fCd23Cre/+
Dot1lf/fCd23Cre/+

Figure S4

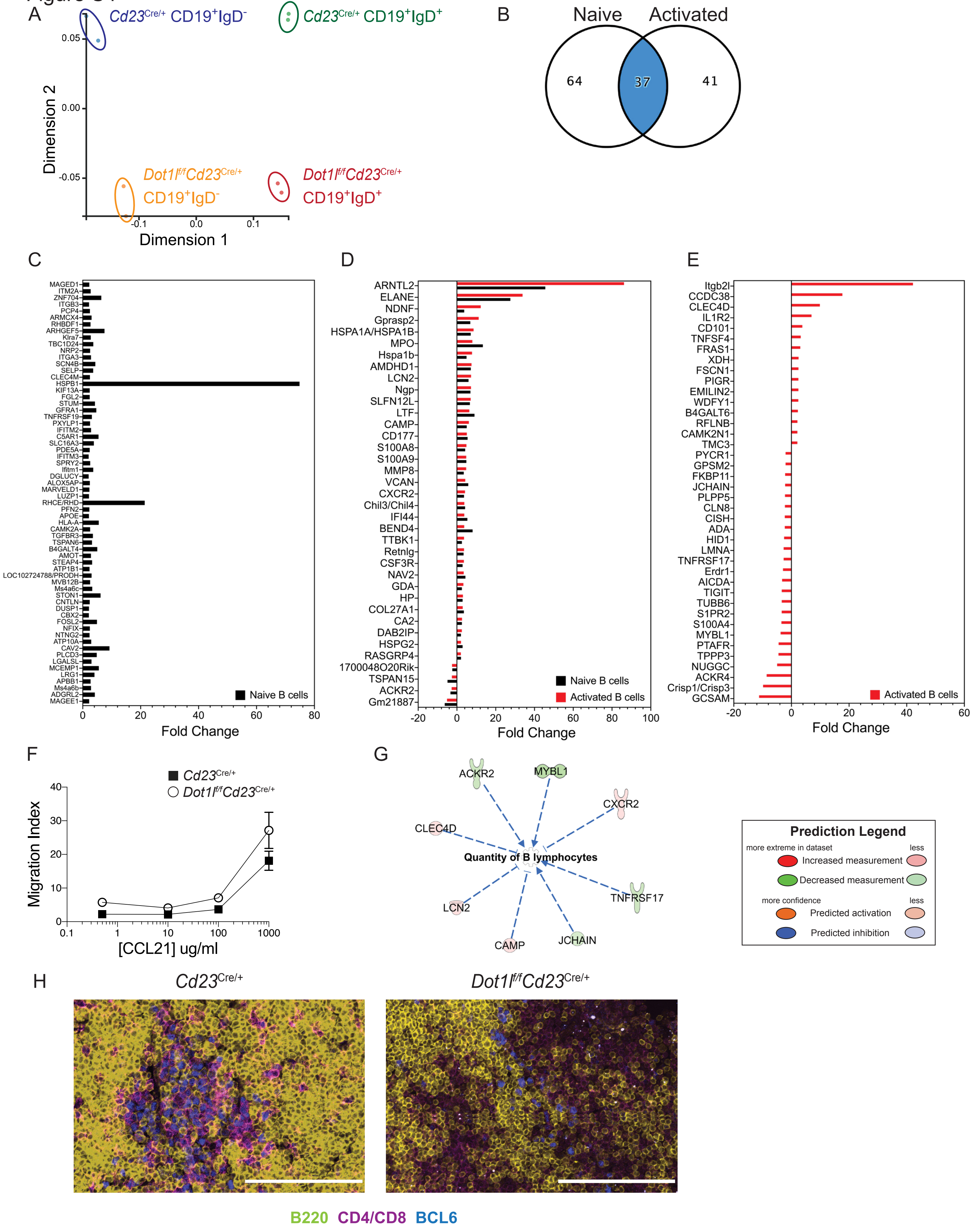
A
Cd23Cre/+ CD19+IgD-
Cd23Cre/+ CD19+IgD+
Dot1lf/fCd23Cre/+ CD19+IgD-
Dot1lf/fCd23Cre/+ CD19+IgD+
Dimension 1
Dimension 2
B
Naive
Activated
64
37
41
C
Naive B cells
Fold Change
D
Naive B cells
Activated B cells
Fold Change
E
Activated B cells
Fold Change
F
Cd23Cre/+
Dot1lf/fCd23Cre/+
Migration Index
[CCL21] ug/ml
G
ACKR2
MYBL1
CXCR2
CLEC4D
Quantity of B lymphocytes
TNFRSF17
LCN2
CAMP
JCHAIN
Prediction Legend
more extreme in dataset
less
Increased measurement
Decreased measurement
more confidence
less
Predicted activation
Predicted inhibition
H
Cd23Cre/+
Dot1lf/fCd23Cre/+
B220 CD4/CD8 BCL6

**SUPPLEMENTAL INFORMATION**

***Figure S1: Conditional Dot1l deletion disrupts B cell development. Related to Figure 1.***

(A) Schematic of conditional deletion of *Dot1l* in the B cell lineage. (B) Representative image of spleens isolated from $Mb1^{Cre/+}$ (left) and $Dot1l^{f/f}$ $Mb1^{Cre/+}$ (right) mice. (C) Spleen weight of naïve, adult $Mb1^{Cre/+}$ (black bars) and $Dot1l^{f/f}Mb1^{Cre/+}$ mice (light grey bars). Results are combined from 2 independent experiments. (D) Flow cytometric analyses of $CD19^{+}$ B cells in the spleen of naïve, adult $Mb1^{Cre/+}$ (black bars), $Dot1l^{fl+}$ $Mb1^{Cre/+}$ (grey bars) and $Dot1l^{f/f}Mb1^{Cre/+}$ mice (light grey bars). (E) Representative images from histological analyses of spleens from naïve mice: MOMA-1 (yellow) and B220 (magenta). Scale bar = 100μm. (F-I) Frequency and number of (F-G) marginal zone B cells and (H-I) follicular B cells in the spleen. (J-O) Different antibody isotypes from the serum of $Mb1^{Cre/+}$ (black squares) and $Dot1l^{f/+}$ and $Dot1l^{f/f}$ $Mb1^{Cre/+}$ mice (white circles) were compared by ELISA analysis: (J) total IgM, (K) total IgG1, (L) total IgG2c, (M) total IgG2b, (N) total IgG3 and (O) total IgA serum levels for each mouse group (n=6). Sigmoidal curve fit of serially diluted samples. (P-Q) Flow cytometric analyses of B cell populations in the bone marrow of naïve, adult $Mb1^{Cre/+}$ (black bars), $Dot1l^{fl+}$ $Mb1^{Cre/+}$ (grey bars) and $Dot1l^{f/f}Mb1^{Cre/+}$ mice (light grey bars). (P) $B220^{+}$ populations, (Q) Pro B cell ($B220^{lo}CD24^{+}BP1^{-}CD43^{+}IgM^{-}$), Pre B cell ($B220^{lo}CD24^{+}BP1^{+}CD43^{-}IgM^{-}IgD^{-}$), Immature B cell ($B220^{+}CD24^{+}CD43^{-}IgM^{lo}IgD^{-}$), Transitional B cell ($B220^{+}CD24^{+}CD43^{-}IgM^{hi}IgD^{-}$), Early Mature B cell ($B220^{+}CD24^{+}CD43^{-}IgM^{hi}IgD^{+}$) and Late Mature B cell ($B220^{+}CD24^{+}CD43^{-}IgM^{lo}IgD^{+}$) populations in the bone marrow. $Mb1^{Cre/+}$ (n=8), $Dot1l^{f/+}Mb1^{Cre/+}$ (n=5) and $Dot1l^{f/f}Mb1^{Cre/+}$ (n=7) mice. Results are combined from four independent experiments. * $P < 0.05$; ** $P < 0.01$; *** $P < 0.001$.

***Table S1: Differential gene expression in CD19$^{+}$IgD$^{+}$ B cells. Related to Figure 3.***

***Table S2: Differential gene expression in CD19$^{+}$IgD$^{-}$ B cells. Related to Figure 3.***

***Table S1: Differential gene expression in CD19$^{+}$IgD$^{+}$ (naïve) B cells. Related to Figure 3.***

| Gene.Name | Naive-WT | Abs fold change Naive-Dot1L | FDR | AveExpr | P value |
|---|---|---|---|---|---|
| Hspa1b | 0 | 2.313857997 | 3.83E-09 | 6.205452252 | 7.51E-13 |
| Hspa1a | 0 | 2.82997678 | 3.83E-09 | 4.879616986 | 3.78E-13 |
| S100a9 | 0 | 2.285418632 | 3.34E-08 | 4.605694001 | 9.83E-12 |
| Ms4a6b | 0 | 1.528215408 | 5.93E-07 | 5.57400116 | 2.33E-10 |
| S100a8 | 0 | 2.092026557 | 7.72E-07 | 3.792594297 | 4.22E-10 |
| Ngp | 0 | 2.804709673 | 7.72E-07 | 2.669183804 | 4.55E-10 |
| Itgb3 | 0 | 1.13041309 | 9.92E-07 | 5.576565487 | 6.81E-10 |
| Ltf | 0 | 3.179306222 | 1.38282E-06 | 2.532179342 | 1.08E-09 |
| Atp1b1 | 0 | 1.252418603 | 2.53446E-06 | 5.392068551 | 2.49E-09 |
| Col27a1 | 0 | 1.859695931 | 3.54786E-06 | 3.353368001 | 3.83E-09 |
| Rhbdf1 | 0 | 1.439365291 | 1.57856E-05 | 4.262392664 | 2.01E-08 |
| 9030617O03Rik | 0 | 1.051158964 | 1.57856E-05 | 4.689808408 | 1.97E-08 |
| Lcn2 | 0 | 2.591904592 | 1.66972E-05 | 1.771857671 | 2.29E-08 |
| Chil3 | 0 | 2.081013798 | 2.09966E-05 | 2.477693241 | 3.09E-08 |
| Ms4a6c | 0 | 1.729239094 | 5.85118E-05 | 4.199634789 | 1.32E-07 |
| Apoe | 0 | 1.0980216 | 7.70096E-05 | 8.22855665 | 2.05E-07 |
| Itm2a | 0 | 1.466180241 | 7.70096E-05 | 3.447344365 | 2.11E-07 |
| Hspg2 | 0 | 1.548911188 | 0.000106919 | 4.070747525 | 3.25E-07 |
| Arntl2 | 0 | 5.511843755 | 0.000181506 | -2.73078596 | 6.48E-07 |
| Mpo | 0 | 3.742536309 | 0.000214381 | -0.24037608 | 7.99E-07 |
| Csf3r | 0 | 1.552258831 | 0.000228252 | 2.32150269 | 8.73E-07 |
| Car2 | 0 | 1.360764256 | 0.000260148 | 3.77172544 | 1.13711E-06 |
| Gm21887 | 0 | -2.65104445 | 0.000315981 | 1.524836693 | 1.45642E-06 |
| Camp | 0 | 2.347657962 | 0.000317715 | 1.018744522 | 1.49557E-06 |
| Alox5ap | 0 | 1.302676948 | 0.000382886 | 2.776062134 | 1.9206E-06 |
| Ackr2 | 0 | -1.73638283 | 0.000656593 | 3.298832147 | 3.79906E-06 |
| Kif13a | 0 | 1.281668271 | 0.00074443 | 3.502716131 | 4.38029E-06 |
| Gm21967 | 0 | -1.92678120 | 0.000906051 | 1.382477832 | 5.81321E-06 |
| Steap4 | 0 | 1.701807073 | 0.000906051 | 2.660848256 | 5.86441E-06 |
| Ttbk1 | 0 | 1.371511953 | 0.000916656 | 1.924829065 | 6.11284E-06 |
| Retnlg | 0 | 1.787442439 | 0.001017787 | 1.632434175 | 6.95379E-06 |
| Cxcr2 | 0 | 1.881454752 | 0.001140607 | 1.198105702 | 8.16557E-06 |
| Hp | 0 | 1.548856427 | 0.001306454 | 1.983894317 | 9.73723E-06 |
| Bend4 | 0 | 3.014315583 | 0.001456727 | 0.446620354 | 1.1143E-05 |
| Cd177 | 0 | 2.462732035 | 0.001650956 | 0.320587863 | 1.39112E-05 |
| Apbb1 | 0 | 1.452560022 | 0.001650956 | 1.979602247 | 1.39239E-05 |
| 1700048O20Rik | 0 | -1.26247913 | 0.001701393 | 3.20130498 | 1.4683E-05 |
| Tnfrsf19 | 0 | 1.658945857 | 0.001707939 | 2.943099025 | 1.5242E-05 |

| | | | | | |
|---|---|---|---|---|---|
| Adgrl2 | 0 | 2.066152913 | 0.00174166 | 2.717190482 | 1.57137E-05 |
| Ifitm3 | 0 | 1.041676312 | 0.001857851 | 2.679912128 | 1.71264E-05 |
| Mcemp1 | 0 | 2.478076436 | 0.001857851 | 0.183256946 | 1.70511E-05 |
| Elane | 0 | 4.785299169 | 0.002172892 | -2.94367441 | 2.21615E-05 |
| Marveld1 | 0 | 1.211773742 | 0.002172892 | 2.314742834 | 2.14306E-05 |
| Dusp1 | 0 | 1.119288778 | 0.002264234 | 6.027346769 | 2.35079E-05 |
| Mmp8 | 0 | 1.825587502 | 0.002264234 | 0.636293431 | 2.35372E-05 |
| Slc16a3 | 0 | 1.943393827 | 0.002346007 | 0.409492658 | 2.5758E-05 |
| Luzp1 | 0 | 1.106681635 | 0.002346007 | 2.483696189 | 2.53382E-05 |
| Atp10a | 0 | 1.555034482 | 0.002394575 | 1.815360667 | 2.67708E-05 |
| Tspan6 | 0 | 1.683211231 | 0.002975673 | 1.081396968 | 3.70609E-05 |
| B4galt4 | 0 | 2.326989425 | 0.003060935 | -0.23441828 | 4.09806E-05 |
| Tbc1d24 | 0 | 1.87470869 | 0.003060935 | 0.834015226 | 4.10647E-05 |
| Cbx2 | 0 | 1.08250632 | 0.003079969 | 2.938635372 | 4.16824E-05 |
| Pcp4 | 0 | 1.320897754 | 0.003107206 | 1.528581734 | 4.26605E-05 |
| Nrp2 | 0 | 1.365526522 | 0.003206457 | 3.75968521 | 4.5281E-05 |
| Rasgrp4 | 0 | 1.088839392 | 0.003988283 | 3.114169848 | 5.93773E-05 |
| Scn4b | 0 | 2.1294652 | 0.003988283 | -0.12733163 | 5.94507E-05 |
| 6330403A02Rik | 0 | 2.080020015 | 0.004101135 | 0.866337063 | 6.23395E-05 |
| Hspb1 | 0 | 6.226554808 | 0.004191745 | -0.56774818 | 6.4539E-05 |
| Cav2 | 0 | 3.208853709 | 0.004325722 | -1.34952917 | 6.82986E-05 |
| Vcan | 0 | 2.554364152 | 0.004531772 | 0.062317339 | 7.20832E-05 |
| Nav2 | 0 | 2.142143535 | 0.00462692 | 1.901764099 | 7.48693E-05 |
| Cd209a | 0 | 1.323944178 | 0.00462692 | 0.611611915 | 7.45903E-05 |
| Cntln | 0 | 1.27887809 | 0.004929706 | 2.136596557 | 8.2524E-05 |
| Ifitm2 | 0 | 1.535979716 | 0.004929706 | 0.82902008 | 8.26694E-05 |
| Itga3 | 0 | 1.529484848 | 0.005228791 | 1.45404272 | 9.28127E-05 |
| Maged1 | 0 | 1.18784316 | 0.005228791 | 2.817588375 | 9.27159E-05 |
| Amot | 0 | 1.513932927 | 0.005254769 | 0.543796854 | 9.37891E-05 |
| H2-Q10 | 0 | 2.470489009 | 0.00565686 | -0.3695914 | 0.000106255 |
| Gfra1 | 0 | 2.22644161 | 0.00565686 | 0.440945447 | 0.000107623 |
| Mvb12b | 0 | 1.359725719 | 0.00565686 | 1.840631278 | 0.000104283 |
| Zfp704 | 0 | 2.678941612 | 0.005774624 | -0.22069523 | 0.00011043 |
| Armcx4 | 0 | 1.614749749 | 0.005789261 | 0.929017064 | 0.000111277 |
| Tspan15 | 0 | -2.27358750 | 0.00580985 | 2.033337497 | 0.000115681 |
| Ifi44 | 0 | 2.430675976 | 0.00580985 | 0.523755385 | 0.00011463 |
| Rhd | 0 | 4.421461976 | 0.00580985 | -1.89216823 | 0.000113425 |
| Magee1 | 0 | 1.162911915 | 0.005939235 | 2.003540417 | 0.000120567 |
| Klra1 | 0 | 1.480750803 | 0.006004832 | 1.193340345 | 0.000122488 |
| Ifitm1 | 0 | 1.864346011 | 0.006216178 | -0.02196145 | 0.000128018 |
| Fgl2 | 0 | 1.274622624 | 0.006622455 | 2.95040722 | 0.000140062 |
| Slfn4 | 0 | 2.741002326 | 0.006774663 | -0.73901729 | 0.000146258 |

| | | | | | |
|---|---|---|---|---|---|
| Nfix | 0 | 1.303990551 | 0.006774663 | 1.245153007 | 0.000147492 |
| Lrg1 | 0 | 2.023849417 | 0.006899636 | 0.242023401 | 0.000150889 |
| Gda | 0 | 1.368179881 | 0.007218036 | 2.344464388 | 0.000164223 |
| Spry2 | 0 | 1.373857297 | 0.007411651 | 1.615658306 | 0.000171536 |
| Dab2ip | 0 | 1.063512918 | 0.007602159 | 3.0441097 | 0.00017669 |
| Lgalsl | 0 | 1.60319526 | 0.00764702 | 1.476208954 | 0.000180733 |
| Camk2a | 0 | 1.360303528 | 0.00764702 | 2.052113313 | 0.000180453 |
| C5ar1 | 0 | 2.447922829 | 0.007694422 | -0.42316542 | 0.000182608 |
| Ndnf | 0 | 1.905522971 | 0.00796378 | -0.52665119 | 0.000194946 |
| Prodh | 0 | 1.615569057 | 0.00811799 | 0.812653497 | 0.000201417 |
| Pde5a | 0 | 1.274629829 | 0.008192762 | 1.409352637 | 0.000206486 |
| Amdhd1 | 0 | 2.83971926 | 0.008240736 | -1.29764159 | 0.000208503 |
| Plcd3 | 0 | 2.282749986 | 0.008803824 | 1.204021593 | 0.000228794 |
| Tgfbr3 | 0 | 1.814568533 | 0.008897631 | 0.297436996 | 0.00023332 |
| Selp | 0 | 1.856638972 | 0.009014327 | -0.36317281 | 0.000240425 |
| Ston1 | 0 | 2.625886405 | 0.009130206 | 0.293250105 | 0.000247564 |
| Pxylp1 | 0 | 1.340622999 | 0.009408159 | 1.350176515 | 0.000258339 |
| Gprasp2 | 0 | 2.786865511 | 0.009537935 | -1.55393697 | 0.000262838 |
| Pfn2 | 0 | 1.208075063 | 0.009556168 | 1.90405161 | 0.000264278 |
| Arhgef5 | 0 | 2.914275509 | 0.009566016 | -1.40633817 | 0.000265488 |
| Ntng2 | 0 | 1.249239098 | 0.009687143 | 2.364384356 | 0.0002698 |
| Fosl2 | 0 | 2.299255193 | 0.009973047 | 1.079428364 | 0.000279389 |

*Table S2: Differential gene expression in CD19$^+$IgD$^-$ B cells. Related to Figure 3.*

| Gene.Name | Act-WT | Abs Fold Change Act-Dot1L | FDR | AveExpr | P value |
|---|---|---|---|---|---|
| Hspa1b | 0 | 2.963011017 | 4.34E-10 | 6.205452252 | 8.06E-14 |
| Hspa1a | 0 | 3.102334636 | 4.34E-10 | 4.879616986 | 8.51E-14 |
| Ackr2 | 0 | -1.450772663 | 1.34E-07 | 3.298832147 | 3.94E-11 |
| S100a9 | 0 | 2.263724375 | 1.51E-07 | 4.605694001 | 5.91E-11 |
| Ngp | 0 | 2.860646155 | 1.52148E-06 | 2.669183804 | 7.46E-10 |
| S100a8 | 0 | 2.296262808 | 2.79486E-06 | 3.792594297 | 1.64E-09 |
| Wdfy1 | 0 | 1.271289152 | 6.0492E-06 | 4.96865679 | 4.15E-09 |
| Ltf | 0 | 2.683125266 | 1.22118E-05 | 2.532179342 | 9.58E-09 |
| Tspan15 | 0 | -1.367851595 | 1.45721E-05 | 2.033337497 | 1.29E-08 |
| Lcn2 | 0 | 2.86545773 | 3.00007E-05 | 1.771857671 | 3.81E-08 |
| Col27a1 | 0 | 1.595863026 | 3.00007E-05 | 3.353368001 | 3.82E-08 |
| Mybl1 | 0 | -1.905282256 | 3.05435E-05 | 1.832856284 | 4.21E-08 |
| Lmna | 0 | -1.432948032 | 3.15025E-05 | 2.69732882 | 5.17E-08 |
| Tppp3 | 0 | -2.167100014 | 3.35676E-05 | 0.863801565 | 6.25E-08 |
| Arntl2 | 0 | 6.43013515 | 4.69155E-05 | -2.73078596 | 9.27E-08 |
| Chil3 | 0 | 1.936405848 | 6.58345E-05 | 2.477693241 | 1.49E-07 |
| 1700048O20Rik | 0 | -1.333227954 | 7.51845E-05 | 3.20130498 | 1.84E-07 |
| Cish | 0 | -1.197333307 | 7.85334E-05 | 1.925283523 | 2.08E-07 |
| Ptafr | 0 | -2.155105866 | 8.03163E-05 | 0.94943737 | 2.28E-07 |
| Fscn1 | 0 | 1.300791748 | 8.03163E-05 | 2.224109896 | 2.22E-07 |
| Nav2 | 0 | 1.811790407 | 8.27242E-05 | 1.901764099 | 2.51E-07 |
| Nuggc | 0 | -2.305016236 | 9.37958E-05 | 1.898574667 | 3.04E-07 |
| Itgb2l | 0 | 5.398933447 | 0.000112991 | -2.71417215 | 3.90E-07 |
| Tnfsf4 | 0 | 1.697711699 | 0.000151561 | 0.336927158 | 5.56E-07 |
| Pigr | 0 | 1.285121108 | 0.000155526 | 2.818614997 | 6.31E-07 |
| Gm21887 | 0 | -2.360590592 | 0.000283807 | 1.524836693 | 1.32649E-06 |
| Csf3r | 0 | 1.860464851 | 0.000365817 | 2.32150269 | 1.90137E-06 |
| Camp | 0 | 2.625197311 | 0.000393741 | 1.018744522 | 2.17399E-06 |
| Aicda | 0 | -1.685516964 | 0.000399143 | -0.16864192 | 2.2703E-06 |
| Fras1 | 0 | 1.622493896 | 0.000401568 | -0.66645710 | 2.32348E-06 |
| Tubb6 | 0 | -1.745082994 | 0.000632179 | 0.682645537 | 3.87853E-06 |
| Osgin1 | 0 | -1.245300231 | 0.000632179 | 2.197293988 | 3.90579E-06 |
| S1pr2 | 0 | -1.756991175 | 0.000804046 | 2.380981776 | 5.04648E-06 |
| Crisp1 | 0 | -3.300227456 | 0.000934408 | -1.12595260 | 6.3799E-06 |
| Hspg2 | 0 | 1.088707641 | 0.000934408 | 4.070747525 | 6.46449E-06 |
| Ppapdc1b | 0 | -1.14898672 | 0.000934408 | 2.62607476 | 6.25076E-06 |
| Gcsam | 0 | -3.490270123 | 0.00101548 | -0.53863487 | 7.36937E-06 |
| Fkbp11 | 0 | -1.119574903 | 0.001126245 | 2.717657325 | 8.50455E-06 |

| | | | | | |
|---|---|---|---|---|---|
| S100a4 | 0 | -1.796461884 | 0.001379105 | 0.723824301 | 1.1211E-05 |
| Cln8 | 0 | -1.163385706 | 0.001532173 | 1.413029855 | 1.33729E-05 |
| Mmp8 | 0 | 2.255963928 | 0.001532173 | 0.636293431 | 1.31669E-05 |
| Ackr4 | 0 | -3.10011899 | 0.001532173 | -1.93229685 | 1.32916E-05 |
| Cxcr2 | 0 | 2.069240736 | 0.001597484 | 1.198105702 | 1.40996E-05 |
| Gm21967 | 0 | -1.956898624 | 0.001822012 | 1.382477832 | 1.73791E-05 |
| Ndnf | 0 | 3.618676879 | 0.001822012 | -0.52665119 | 1.75108E-05 |
| Ccdc38 | 0 | 4.144568256 | 0.001868973 | -1.72777374 | 1.83287E-05 |
| Elane | 0 | 5.082166094 | 0.001955334 | -2.94367441 | 1.99426E-05 |
| Gprasp2 | 0 | 3.490883479 | 0.001955334 | -1.55393697 | 1.97794E-05 |
| Fam101b | 0 | 1.069817226 | 0.002037602 | 5.587118514 | 2.20299E-05 |
| Mpo | 0 | 2.997407607 | 0.002081724 | -0.24037608 | 2.3069E-05 |
| Ada | 0 | -1.206160768 | 0.002158388 | 3.173973269 | 2.76771E-05 |
| Car2 | 0 | 1.410336274 | 0.002158388 | 3.77172544 | 2.62527E-05 |
| Ifi44 | 0 | 1.920036673 | 0.002158388 | 0.523755385 | 2.76128E-05 |
| Camk2n1 | 0 | 1.04577939 | 0.002158388 | 1.04838751 | 2.59448E-05 |
| Ttbk1 | 0 | 1.889412322 | 0.002343803 | 1.924829065 | 3.03405E-05 |
| B4galt6 | 0 | 1.169611758 | 0.00243601 | 0.540855937 | 3.24233E-05 |
| Jchain | 0 | -1.130510644 | 0.002999923 | 7.83805826 | 4.38353E-05 |
| Hp | 0 | 1.643155577 | 0.003035657 | 1.983894317 | 4.49369E-05 |
| Hid1 | 0 | -1.340404776 | 0.003359416 | 1.402147505 | 5.17239E-05 |
| Amdhd1 | 0 | 2.907492533 | 0.003553253 | -1.29764159 | 5.57537E-05 |
| Retnlg | 0 | 1.882665812 | 0.003690474 | 1.632434175 | 5.9347E-05 |
| Erdr1 | 0 | -1.541826294 | 0.003742839 | 1.52555859 | 6.05637E-05 |
| Cd177 | 0 | 2.351722309 | 0.004053177 | 0.320587863 | 6.71753E-05 |
| Il1r2 | 0 | 2.801296244 | 0.004134054 | 0.039072859 | 6.93266E-05 |
| Bend4 | 0 | 1.910190155 | 0.004348129 | 0.446620354 | 7.63278E-05 |
| Gpsm2 | 0 | -1.027515451 | 0.004644209 | 0.991933497 | 8.39913E-05 |
| Gda | 0 | 1.729824062 | 0.004762996 | 2.344464388 | 8.73473E-05 |
| Slfn4 | 0 | 2.82326356 | 0.005472365 | -0.73901729 | 0.00010626 |
| Xdh | 0 | 1.318491944 | 0.005698344 | 1.951240169 | 0.000111206 |
| Pycr1 | 0 | -1.025345269 | 0.0059004 | 1.533510018 | 0.000119779 |
| Tnfrsf17 | 0 | -1.437967394 | 0.006140709 | -0.90241905 | 0.000127737 |
| Dab2ip | 0 | 1.342662908 | 0.006140709 | 3.0441097 | 0.00012634 |
| Clec4d | 0 | 3.303470614 | 0.006557919 | -0.40873811 | 0.000139558 |
| Vcan | 0 | 2.099137248 | 0.007772043 | 0.062317339 | 0.00018445 |
| Tigit | 0 | -1.743921519 | 0.00853285 | -0.10225920 | 0.000211932 |
| Cd101 | 0 | 1.923702091 | 0.008766411 | 0.65004755 | 0.000224383 |
| Emilin2 | 0 | 1.28263613 | 0.009539102 | 1.504554161 | 0.000250709 |
| Rasgrp4 | 0 | 1.026352151 | 0.009909572 | 3.114169848 | 0.000273079 |
| Tmc3 | 0 | 1.042977263 | 0.009909572 | -0.11885847 | 0.000270808 |
